# Fueling a predator death-trap: Trophic subsidies and the risk of management-induced collapse in a predator-prey system

**DOI:** 10.1101/2025.08.26.672345

**Authors:** Vinicius P. O. Gasparotto, Tatiane Micheletti, Paulo R. Mangini, Jean C. R. Silva, Ricardo A. Dias

## Abstract

1. Anthropogenic subsidies create complex ecological dynamics, yet their interaction with invasive species management is poorly understood. Managing subsidies or predators in isolation risks perverse outcomes, including population collapses, demanding a more holistic understanding.
2. We employed a Pattern-Oriented Inference framework to synthesize multiple lines of evidence for the endemic Noronha skink (*Trachylepis atlantica*) across an archipelago. We analyzed three key patterns using complementary methods: (i) population density, estimated via capture-mark-recapture and negative binomial GLMs of count data; (ii) individual body size, using gamma GLMs; and (iii) injury rates, via tail-autotomy analysis.
3. We documented a dramatic density gradient driven by predation pressure. Skink populations were over three times denser on predator-light secondary islands (0.411 ind/m^2^, 95% CI: 0.297–0.568) than in predator-rich PARNAMAR (0.125 ind/m^2^, 95% CI: 0.090–0.174). Although anthropogenic subsidies boosted density by 82% in the inhabited APA (0.227 ind/m^2^, 95% CI: 0.174–0.297), this was insufficient to overcome the severe impact of invasive predators.
4. This created a paradox where density, a traditional population success metric, was inverse to size, an indicator of individual fitness. On the main island, although the subsidized APA was denser than PARNAMAR, individual condition was no better. Adult males in the APA were as small as those in the lowest-density sites and significantly smaller than males on the secondary islands. Predation pressure explains this gradient, with tail-loss injury rates peaking in the APA (29.3%), remaining high in PARNA (25.0%), and dropping about 50% on the secondary islands (15.0%).
5. Synthesis and applications: The convergence of these patterns supports a “predator death-trap” hypothesis, where trophic subsidies in the inhabited APA fuel high skink recruitment, masking extreme mortality from a subsidized predator community and other anthropogenic threats. This dynamic produces high population turnover and truncates size structure by selectively removing larger, older adults and may explain patterns seen elsewhere. Our findings have critical management implications: removing food subsidies without concurrent, effective control of key invasive predators could trigger a population collapse. We advocate integrated, multi-species management and provide robust evidence for the threatened status of this endemic reptile.

## 1. INTRODUCTION

Due to their geographic isolation, oceanic islands foster unique evolutionary dynamics, resulting in high rates of endemism and unique community structures (Kier et al., 2009). However, this isolation also makes island biota exceptionally vulnerable to anthropogenic disturbances and biological invasions, primary drivers of the ongoing biodiversity crisis and have caused numerous extinctions (Spatz et al., 2017). Understanding the complex and often interacting impacts of these cumulative pressures is a central challenge for applied ecology (Maeda et al., 2019; Oppel et al., 2014).

The archipelago of Fernando de Noronha, Brazil, exemplifies this challenge. Noronha is home to the endemic Noronha skink (*Trachylepis atlantica*), a lizard that evolved in a historically predator-free environment (Rocha et al., 2009), and an important pollinator species for the native mulungu tree (*Erythrina velutina*). The main island of the archipelago is now inhabited and hosts a rich suite of invasive species, including black rats (*Rattus rattus*), stray and feral cats (*Felis catus*), tegu lizards (*Salvator merianae*), and the cattle egret (*Bubulcus ibis*), all of which are documented predators of the Noronha skink (Gaiotto et al., 2020; Gasparini et al., 2007; Micheletti et al., 2020). In contrast, the smaller, uninhabited secondary islands, while still harboring invasive rats—except for one island, where eradication has been successful—are largely free from cats and tegu lizards (Abrahão et al., 2019; Russell et al., 2018).

Complicating this dynamic is the direct and indirect influence of human settlement on the main island, which is administered as both a protected Environmental Protection Area (APA) and a more strictly protected National Park (PARNAMAR). While both regions host the full suite of invasive predators, the APA, which contains the urban areas, provides anthropogenic food subsidies (e.g., refuse) that can alter resource availability for both skinks and predators (Gasparini et al., 2007). These potential benefits, however, are coupled with a suite of concentrated negative pressures unique to the APA, including direct persecution linked to tourism-related concerns (e.g., business owners fearing that lizards may disturb visitors), alongside predation from domestic and stray animals, and profound habitat modification. This creates a classic management dilemma: decisions must be made in a complex system where such countervailing effects are difficult to separate, but obtaining enough data to inform all parameters is logistically infeasible. For example, although a general decline in skink density on the main island has been noted (AEMA, 2017), the lack of a clear understanding of the relative importance of these interacting drivers has impeded the development of robust conservation strategies and has hindered efforts to formally classify the species as ‘Vulnerable’ or ‘Threatened’ under IUCN criteria (IUCN, 2014).

This archipelago-wide gradient of invasive predator presence, coupled with differences in human influence between management zones, creates a natural experiment for assessing their cumulative impacts, which we use to address this challenge directly. We adopt a Pattern-Oriented Inference (POI) framework —a novel approach for synthesizing the types of disparate datasets common in applied ecology— inspired by the Pattern-Oriented Modelling approach (Grimm et al., 2005), which uses the principle that an explanatory hypothesis gains substantial credibility if it can simultaneously explain multiple, independent patterns observed at different scales or levels of organization. By integrating multiple lines of evidence and focusing on a suite of robust patterns— (1) population density, (2) individual body size, and (3) rates of injury— we test the overarching hypothesis that the interplay between invasive predators and human activities transforms inhabited areas into a ‘predator death-trap,’ which in turn generates a high-turnover population dynamic that functions as an ‘invisible sink’. Our goal is to demonstrate that even with inherent data limitations, a structured synthesis can provide a clear understanding of system dynamics and generate robust recommendations to guide urgent management decisions while avoiding perverse outcomes, such as intensifying predation pressure and causing a collapse of the Noronha skink population. Ultimately, such a framework can provide a strong evidence foundation for the effective wildlife management.

## 2. METHODS

To untangle the complex interactions between invasive species, human activities, and skink populations, we adopted a Pattern-Oriented Inference (POI) framework. This approach, inspired by Pattern-Oriented Modeling (Grimm et al., 1996, 2005; Grimm & Railsback, 2012), uses multiple, independent patterns observed in a system to make robust inferences about underlying ecological processes, even in the face of uncertainty. We focused on three key patterns: population density, individual body size, and non-lethal injury rates, which were analyzed across three distinct management zones within the archipelago, representing a gradient of invasive species pressure and human influence.

### 2.1 Study Area and Design

The study was conducted in the Fernando de Noronha archipelago (3°51’13.71”S, 32°25′25.63”W), a Brazilian federal territory located in the Atlantic Ocean (Figure 1). The archipelago encompasses a total terrestrial area of 18.22 km^2^, dominated by the main island (16.89 km^2^) and a series of smaller secondary islands (1.33 km^2^). The climate of Fernando de Noronha is consistently warm, with sea surface temperatures averaging 27 °C and atmospheric temperatures generally between 25 and 31 °C. Annual precipitation is approximately 1,400 mm, falling predominantly from January through July, while August to December constitutes the drier period (WeatherSpark, 2024). Original vegetation on the archipelago is classified as Seasonal Deciduous Forest, showing marked contrasts between the wet and dry seasons (Teixeira & Linsker, 2003).

**Figure 1.**
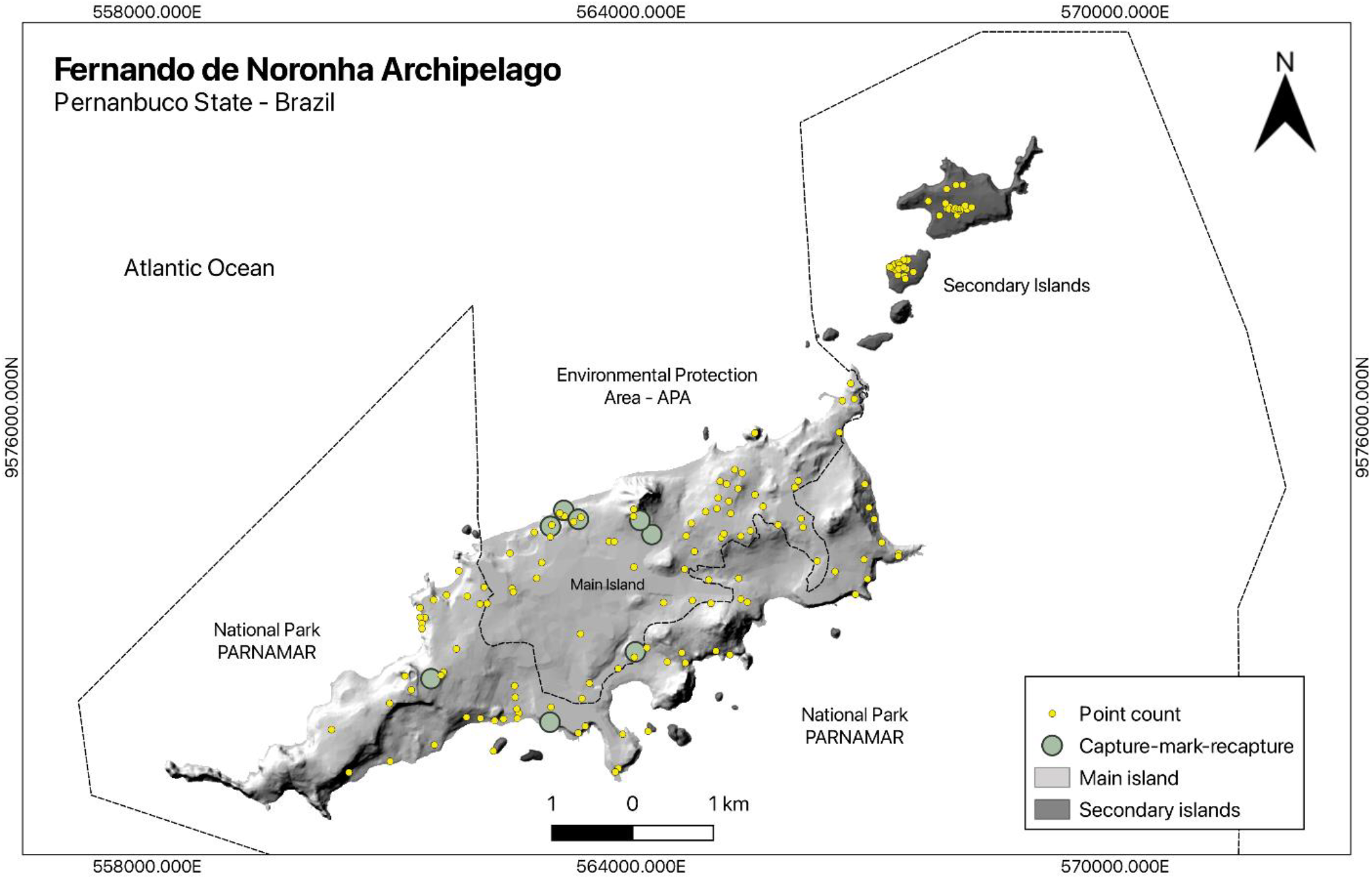
Map of the Fernando de Noronha archipelago, Brazil, depicting the spatial distribution of study sites and sampling efforts in relation to key environmental drivers. The main island is divided into the protected PARNAMAR and the inhabited APA, representing zones with varying anthropogenic subsidies. Secondary islands (dark grey) represent areas with reduced invasive predator presence. Yellow circles mark point count survey locations, and green circles denote capture-mark-recapture survey locations.

The main island—the only inhabited island—is divided into two contiguous federal protected areas that form the basis of our study design. The first is PARNAMAR, the strictly protected Marine National Park (IUCN Category II), which covers approximately 70% of the island’s terrestrial area (11,82 km^2^). This site is uninhabited, lacks direct anthropogenic subsidies, but contains the full suite of key invasive predators: cats (*Felis catus*), tegus, and rats, as well as egrets. The second is the APA, the inhabited Environmental Protection Area (IUCN Category V), which covers the remaining 30% of the island (5,07 km^2^). In 2017, this site supported a resident population of approximately 2,900 people and attracted significant tourism. Like PARNAMAR, it contains the full predator suite with a higher free ranging cat density (Dias et al., 2017), but it is additionally characterized by substantial human presence, including urban development, habitat modification, and the availability of anthropogenic subsidies. A third site was defined to represent a “baseline” environment. These are the ‘secondary islands’—a group of small, uninhabited islands that are largely free of cattle egrets, invasive cats and tegu lizards. While all except one (*Ilha do Meio*) also host invasive black rats, all secondary islands lack the direct human pressure and subsidies found in the APA.

### 2.2 Data Collection and Analysis

To investigate our three core patterns, we employed distinct analytical approaches tailored to the available data for each pattern. All statistical analyses were performed in R version 4.4.2 (R Core Team, 2024). The full workflow including all data and code—models, parameterization, outputs, and diagnostics—are freely available on GitHub (see Data Availability Statement) and provided as an annex to this manuscript (Appendix A).

#### Pattern 1: Population Density

We estimated skink density on the three sites using two complementary methods. First, we used point count data collected at standardized survey locations with a 2-meter radius (area ≈ 12.57 m^2^) to model variation in skink density on PARNAMAR, APA and secondary islands. We fitted a negative binomial generalized linear model (GLM) using the MASS package in R (Venables & Ripley, 2002), with skink counts as the response variable. The model included two fixed-effect predictors: ‘invasive species presence’ (*InvasiveSpeciesPresence*), a binary factor indicating the presence of the full invasive predator suite versus a reduced suite on the secondary islands, and ‘food supplementation’ (*FoodSupplementation*), a binary factor representing the presence or absence of anthropogenic subsidies. To account for the size of the survey area and express the results in terms of density, we included the natural logarithm of the survey area as an offset term. We explored two alternative model formulations to test the robustness of our results. These included (i) a model with an interaction term between ‘invasive species presence’ and ‘food supplementation’, and (ii) a generalized linear mixed-effects model (GLMM) with a random intercept for study site to account for potential spatial grouping. Model comparison using Akaike Information Criterion (AIC) confirmed that the simpler GLM provided the best fit to the data, as the interaction term was not estimable due to data sparseness and the random intercept in the GLMM showed negligible variance. We therefore selected the main effects GLM for inference and assessed its adequacy using simulation-based residual diagnostics via the DHARMa package in R (Hartig, 2024), which included checks for overdispersion, zero-inflation, and uniformity in residual distributions.

Second, for a more detailed assessment of skink density on the main island, we conducted a capture-mark-recapture analysis within the APA and PARNAMAR sites. We applied a Poisson-log normal mark-resight model using the RMark package in R (Laake, 2013), which allowed joint estimation of the resighting probability (*alpha*) and the super-population size (*U*) at each site. We constructed a candidate set of five biologically motivated models to test the influence of site and sampling effort on detection and population size parameters. Models were ranked based on AIC corrected for small sample sizes (AICc), and the model with the lowest AICc was selected for inference.

#### Pattern 2: Individual Body Size

To investigate impacts on individuals body size, we analyzed head length as a stable morphometric trait. Unlike variables such as tail length or body mass, which may vary due to injury or recent feeding, head length reflects skeletal structure and is not affected by tail loss or transient weight changes. It also provides a more reliable and consistent measure than snout–vent length, which can be more error-prone due to curvature or movement during handling. Given that head length is a continuous and strictly positive variable, we fitted a gamma generalized linear model (GLM) with a log link function. An initial model tested for main effects of sex and site on head length. To assess whether environmental conditions affected males and females differently, a second model included an interaction term (sex × site). Model fit was evaluated using diagnostic plots, including residuals versus fitted values and Q-Q plots, to verify assumptions of the gamma distribution. Estimated marginal means and post-hoc pairwise comparisons among group levels were performed with Tukey-adjusted p-values were computed using the emmeans package (Lenth, 2025).

#### Pattern 3: Injury Rates (signs of tail autotomy)

To assess variation in predation pressure across regions, we quantified the frequency of tail autotomy (i.e., tail loss or signs of regrowth). While intraspecific aggression is a known cause of tail autotomy in this species (Gasparotto, 2021), a difference in prevalence of injury across sites can serve as a robust proxy for sublethal predation pressure. For each individual, the presence or absence of autotomy was recorded, and a contingency table was constructed to summarize the number of injured versus uninjured individuals across the three sites. We then used Pearson’s Chi-squared test, implemented in base R (R Core Team, 2024), to test whether the proportion of individuals with autotomy differed significantly among sites. Given the potential for low statistical power due to limited sample sizes, we conducted a detailed post-hoc power analysis to formally evaluate the sensitivity of our Chi-squared test. This analysis, using the pwr package in R (Champely, 2020), included: (i) calculating the achieved power of the 3×2 omnibus test based on the observed effect size (Cohen’s w); (ii) estimating the total sample size required to achieve 80% power; (iii) calculating pairwise power for two-proportion tests with unequal sample sizes; and (iv) determining the minimum detectable difference (MDD) at 80% power for key contrasts involving the smallest sample sizes. To validate these analytic results, we also performed a simulation-based power analysis with 5,000 replicates.

## 3. RESULTS

Our analyses revealed three distinct patterns in the Noronha skink populations converging to one theory: the complex interplay between invasive predators and anthropogenic subsidies.

### Pattern 1: A Density Gradient Driven by Invasive Predators and Food Subsidies

The skink population density varied significantly across the three sites. For the negative binomial generalized linear model of point counts, residual diagnostics confirmed a good model fit, with simulated residuals closely following expected distributions and no evidence of systematic deviation, overdispersion, or outliers. The model explained a modest but reasonable proportion of the variance (Nagelkerke’s R^2^ = 0.17), as is typical for ecological count data. The model estimated the highest density on the secondary islands (0.411 individuals/m^2^, 95% CI: 0.295–0.573) and significantly lower densities on the predator-rich main island, with the subsidized APA (0.204 ind/m^2^, 95% CI: 0.153–0.273) having a higher density than the unsubsidized PARNAMAR (0.156 ind/m^2^, 95% CI: 0.115–0.213), although not statistically significant (Figure 2). This pattern was, however, further clarified by the more detailed capture-mark-recapture analysis.

**Figure 2.**
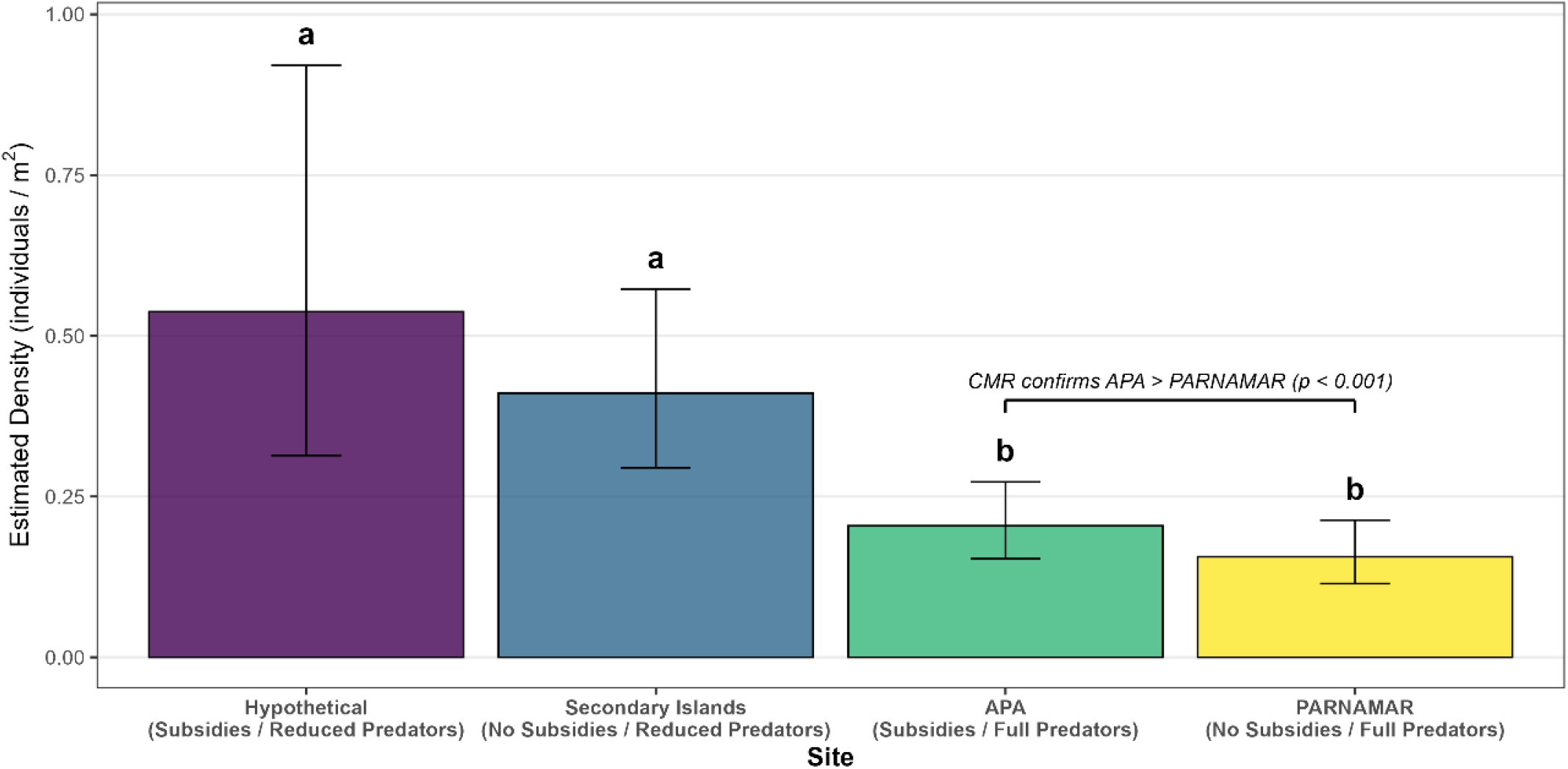
Skink density is strongly suppressed by the full invasive predator suite, a conclusion refined by a capture-mark-recapture analysis. The main bars show estimated marginal mean densities of the Noronha skink (individuals /m^2^) from a Negative Binomial GLM of broad-scale point counts. Error bars represent 95% confidence intervals. Different letters (a, b) indicate statistically significant differences based on this GLM, which shows that sites with a reduced predator suite support significantly higher densities than main island sites. The bracket and annotation highlight a key finding from a separate, more detailed capture-mark-recapture analysis conducted only on the main island: skink population size was confirmed to be significantly higher in the subsidized APA compared to the unsubsidized PARNAMAR (p < 0.001).

The top-ranked capture-mark-recapture model, which included an effect of site on super-population size (*U*), had the lowest AICc (855.39; ΔAICc = 1.80 compared to the next best model; Appendix A, p. 40). Estimated population sizes were significantly lower in PARNAMAR colonies compared to those in the APA (β = −2.00, p < 0.001), despite a significantly higher observation probability for PARNAMAR (β = 0.41, p < 0.05). Together, these results establish a clear pattern, where secondary islands present a significantly larger population density as APA, which in turn, presents a larger population than PARNAMAR.

### Pattern 2: Size Structure Reveals Demographic Truncation on the Main Island

The population’s size structure did not directly correspond to the density pattern, revealing a critical discrepancy on the main island. While the APA supported a higher density than PARNAMAR, its adult males were no larger, pointing to differential mortality pressures (Figure 3). A gamma generalized linear model that included an interaction between sex and site provided a better fit to the head length data (AIC = 24.116) than a model with only main effects (AIC = 25.051). The sex × site interaction term was significant (sexM:sitePARNA, p = 0.0285), indicating that differences in size between sites were primarily driven by changes in the male population. Model diagnostics confirmed data quality with no outliers detected, and residuals displayed no major deviations from model assumptions (see Appendix A).

**Figure 3.**
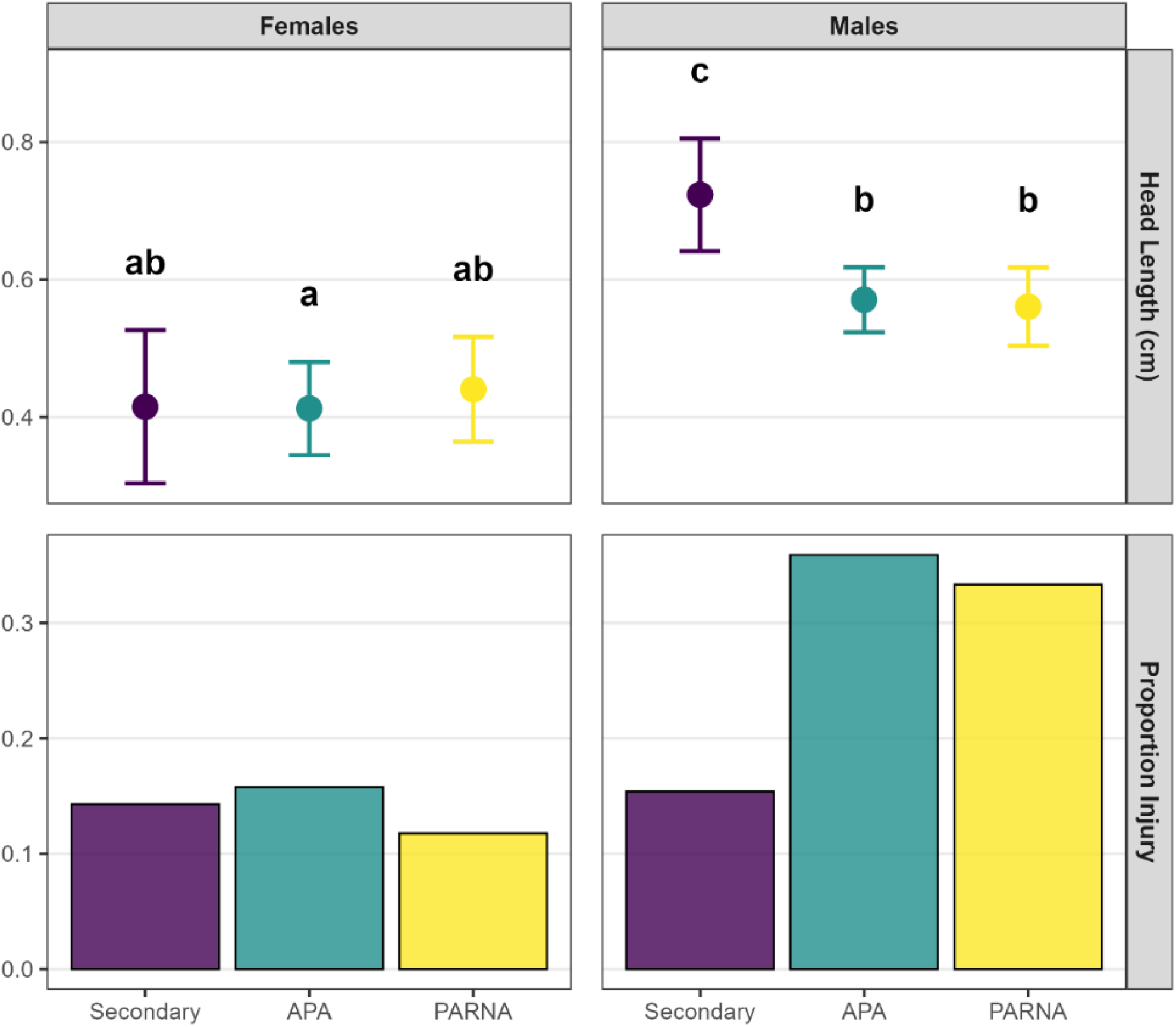
Demographic truncation in male skinks is correlated with higher rates of non-lethal injury. While female skinks (left panels) show no significant differences in head length across sites (mean ± 95% CI), males (right panels) are significantly larger on the secondary islands compared to the main island sites (APA and PARNA) where we observe lower invasive species and anthropogenic pressure. This smaller male body size on the main island coincides with a higher proportion of individuals exhibiting tail autotomy (a proxy for predation pressure), particularly in the APA. Letters indicate significant differences (p < 0.05) in head length only.

Post-hoc pairwise comparisons (Tukey-adjusted) revealed the specific nature of this interaction (Figure 3). While there were no significant size differences among females across the three sites (e.g., females on secondary islands vs. females in the APA, p = 1.00; females in the APA vs. females in the PARNAMAR, p = 0.99), the effect on males was pronounced. Males on the secondary islands were significantly larger than males from both the APA (p = 0.0214) and PARNAMAR (p = 0.0195). There was no statistical size difference between males inhabiting the APA and PARNAMAR (p = 0.9998). This establishes a clear pattern of demographic truncation, where the largest size classes of males are notably absent from the main island populations, regardless of subsidy presence.

### Pattern 3: Injury Rates Align with a High-Pressure Environment on the Main Island

Providing a mechanistic link to the observed size truncation, rates of non-lethal injury (tail autotomy) followed a trend of elevated pressure on the main island. The proportion of injured individuals was highest in the APA (29.3%), followed by PARNAMAR (25.0%), and was lowest on the Secondary Islands (15.0%). Despite this clear directional pattern, the overall difference was not statistically significant (Pearson’s Chi-squared test: χ^2^ = 1.61, df = 2, p = 0.45). Our post-hoc power analysis revealed that this was a consequence of low statistical power. With our realized sample sizes, the achieved power to detect an effect of the observed magnitude was only ∼19%, and an estimated ∼729 individuals would be required to reach the standard 80% power threshold.

The minimum detectable difference for comparisons involving the small secondary island sample was 32.6–34.1 percentage points, indicating only very large effects could be reliably detected. Together, these analyses confirm our interpretation that the non-significant test result is due to statistical limitations rather than the absence of a biological effect. The observed trend of higher injury rates on the main island therefore provides a final, corroborating line of evidence that aligns with the body size data and supports the hypothesis that skinks, particularly in the APA, experience a higher rate of sublethal predator encounters.

## 4. DISCUSSION

Making robust management decisions for species of conservation concern often requires synthesizing information from multiple, imperfect sources of data. In this study, we employed a ‘Pattern-Oriented Inference’ framework to integrate results from population density models, individual morphometrics, and rates of injury to build a cohesive understanding of the complex pressures facing *Trachylepis atlantica*, the island endemic Noronha skink lizard. The convergence of these distinct patterns provides a powerful line of evidence that transcends the limitations of any single analysis. It resolves an apparent ecological paradox by revealing that the inhabited area of the main island functions as a predator death-trap, a dynamic with profound and urgent implications for the conservation of this endemic species.

Our results establish a clear density gradient across the sites (secondary islands > APA > PARNAMAR), which can be explained by the interplay of top-down and bottom-up forces. The high density on the secondary islands serves as a crucial baseline, demonstrating the species’ potential in an environment with reduced predator pressure, even without anthropogenic subsidies. Conversely, the significantly suppressed density in the predator-rich PARNAMAR reflects the strong top-down control exerted by the full suite of invasive predators, particularly cats, tegus, cattle egrets, and rats (Gaiotto et al., 2020; Micheletti et al., 2020). The most insightful finding, however, is the intermediate density in the APA. Here, anthropogenic food subsidies appear to provide a bottom-up stimulus, likely boosting skink recruitment and survival enough to elevate the population density above that of the unsubsidized PARNAMAR. However, while subsidy-driven increases have been observed elsewhere (Plaza & Lambertucci, 2017), our work demonstrates a critical caveat: this apparent recovery can conceal the transition of a habitat into a high-mortality sink.

The key to deciphering the system lies in how size-selective predation resolves the apparent paradox on the main island. Predation is rarely random, and traits that increase conspicuousness, such as large body size and territorial behaviour, often elevate mortality risk. For instance, Bateman and Fleming (2011) demonstrated that large, territorial adult male anoles suffered significantly higher rates of sublethal predation attempts from cats, providing a direct, evidence-based mechanism for the size-selective pressure we propose is acting on the Noronha skink. In the APA, this pressure is amplified because anthropogenic subsidies support high densities of both skinks and their invasive predators (Dias et al., 2017), potentially increasing lethal encounters. Moreover, human activity in the APA reduces the availability of alternative native prey (particularly birds), likely concentrating predation pressure on skinks as one of the few abundant resources remaining. The outcome is a stark pattern of demographic truncation, where the selective removal of the largest and oldest males leaves a high-turnover population dominated by smaller, younger individuals. This directly explains why the subsidized APA, despite supporting more skinks, has males no larger than those in the resource-poor, predator-rich PARNAMAR. The trend of higher injury rates in the APA provides a final, corroborating line of evidence, confirming that this truncated size structure is a consequence of relentless mortality, not stunted growth.

The convergence of these patterns—subsidized density, a truncated male size structure, and elevated injury rates—strongly supports our central hypothesis: the inhabited APA functions as a predator death-trap. Ecological traps occur when an environmental cue, which normally indicates high-quality habitat, leads an organism to a low-quality patch where its fitness is compromised (Patten & Kelly, 2010). In Noronha, the abundance of food in the APA likely serves as an attractive cue for skinks, concentrating their population. However, these same subsidies also support a high-density, subsidized predator community (Dias et al., 2017). The result is a fatal combination of high prey density and high predator density in the same location, creating an intensified predator-prey dynamic that transforms the APA from a subsidized refuge into a population sink.

Our findings reveal a critical and non-obvious management trap that could have catastrophic consequences: removing food subsidies in the APA without first reducing invasive predators. Because subsidies currently fuel high recruitment, eliminating them in isolation would suppress prey production while leaving a dense predator community intact—thereby increasing per-capita predation risk and plausibly precipitating a population collapse, as suggested for the species in theoretical studies (Gasparini et al., 2007). Effective conservation therefore requires an integrated sequence: (i) implement coordinated, sustained control of key invasive predators (e.g., cats, tegus, cattle egrets, rats); (ii) only then phase down anthropogenic subsidies via improved waste management and feeding restrictions; and (iii) track explicit triggers (density, size-structure, and injury rates) under a real-time adaptive monitoring framework (Micheletti et al., 2025). This strategy addresses both top-down and bottom-up drivers simultaneously, avoiding management-induced feedbacks and aligning actions with the multi-line evidence that the APA currently functions as a predator death-trap.

Our integrated results provide strong, cohesive evidence that the Noronha skink is facing cryptic but severe threats that warrants re-assessing the species as at least “Vulnerable” under IUCN criteria (A2: observed population decline; C1: small and declining population size; IUCN, 2014), a conclusion previously obscured by its apparent abundance in the APA. This concern is further heightened by its insular life-history strategy: the species exhibits a reduced, seasonal reproductive pattern compared with continental congeners (Migliore et al., 2017), reflecting an evolutionary history in a predator-free environment. Such traits likely lower its resilience to the novel and intense predation pressures it currently faces. This case study therefore serves as a critical cautionary tale for conservation, demonstrating the danger of relying on single metrics like population density in complex, subsidized ecosystems where a high density can mask an underlying population sink and create a dangerous illusion of security. These findings also inform Brazil’s invasive species strategy, highlighting the need for integrated subsidy and predator control rather than piecemeal actions. Integrated eradication campaigns on islands elsewhere, such as New Zealand (Griffiths et al., 2012), demonstrate the feasibility of coordinated invasive species and subsidies’ control (even if for public-safety or operational control), which could provide a model for Noronha.

Our findings likely extend beyond Noronha. Similar subsidy–predator interactions have been reported for large lizards (Jessop et al., 2012) and free-ranging cats (Fleming et al., 2022; Kazato et al., 2020; Maeda et al., 2019) on continental areas. Food waste has even been directly linked to elevating carrying capacity and sustaining predator guilds (Almaraz et al., 2022). We suggest that the predator death-trap dynamic may be a general risk wherever subsidies elevate recruitment in prey populations but simultaneously fuel predator communities. We demonstrate that a Pattern-Oriented Inference framework provides the necessary tool for managers to synthesize multiple data types—even those with limitations—to uncover these hidden dynamics and form a robust basis for policy. Ultimately, our findings deliver an urgent warning for managers of human-altered ecosystems globally: an integrated management plan that correctly identifies and prioritizes the primary threatening process, in this case invasive predators, is essential to avoid perverse outcomes, such as triggering a preventable population collapse.

