## Appendix A for "Fueling a predator death-trap: Trophic subsidies and the risk of management-induced collapse in a predator-prey system"

### Fueling a predator death-trap: data analysis

Anonymous The Scientist

2025-08-26

### Introduction

This analysis examines Mabuya (Trachylepis atlantica; skink) populations across three distinct sites (i.e., Secondary Islands, PARNAMAR and APA) on Fernando de Noronha archipelago, investigating how invasive species presence and food supplementation affect population density, body size, and injury rates.

#### Study Area Context

The study encompasses the following areas:

- **Main Island: 16.89 km²**
  - PARNAMAR: 11.424 km²
  - APA: 5.466 km²
- **Secondary Islands: 1.33 km²**
- **TOTAL PARNA: 12.754 km²**
- **TOTAL TERRESTRIAL AREA (PARNA + APA): 18.22 km²**

### Package Setup

We'll use the `Require` package for better dependency management across all analyses.

```
if(!require("Require")){
  install.packages("Require")
}
library("Require")

Require::Require("data.table")
Require::Require("emmeans")
Require::Require("DHARMa")
Require::Require("glmmTMB")
Require::Require("performance")
Require::Require("RMark")
Require::Require("xlsx")
Require::Require("ggplot2")
Require::Require("multcompView")
Require::Require("dplyr")
Require::Require("pwr")
```

### Survey Effort Analysis

Before analyzing the biological data, we assesed whether our sampling effort was balanced across the study area.

#### Loading Density Data

```
mabuia_dd <- fread("data/Density_Data.csv")
```

#### Calculating Sampling Effort Distribution

```
pointsCounts <- mabuia_dd[, .N, by = "Site"]
pointsCounts[Site == "APA", pointsArea := 5.466]
pointsCounts[Site == "PARNA", pointsArea := 11.424]
pointsCounts[Site == "Secundaria", pointsArea := 1.33]
pointsCounts[, pointsDensity := N/pointsArea]
pointsCounts[, propPoints := N/(sum(pointsCounts[, N]))]
pointsCounts[, propSize := pointsArea/(sum(pointsCounts[, pointsArea]))]

print("Survey effort distribution:")
```

```
## [1] "Survey effort distribution:"
```

```
print(pointsCounts)
```

```
## Index: <Site>
##          Site     N pointsArea pointsDensity propPoints   propSize
##        <char> <int>      <num>         <num>      <num>      <num>
## 1:        APA    58      5.466     10.611050  0.3866667 0.30000000
## 2:      PARNA    55     11.424      4.814426  0.3666667 0.62700329
## 3: Secundaria    37      1.330     27.819549  0.2466667 0.07299671
```

#### Survey Effort Assessment

There is indeed a bit of unbalance but it is not as critical as it seems:

- **APA**: 39% points over 30% area
- **PARNA**: 37% points over 62% area

- **Secondary**: 25% points over 7% area

The sampling is reasonably proportional to area, with only modest over-sampling of Secondary Islands and slight under-sampling of PARNAMAR relative to area.

### Population Density Analysis

#### Step 1: Survey Area Calculation

Each survey point was a circle with 2m radius, giving us a standardized survey area.

```
mabuia_dd[, survey_area := pi * (2^2)]
print(paste("Survey area per point:", round(mabuia_dd$survey_area[1], 3), "m²"))
```

```
## [1] "Survey area per point: 12.566 m²"
```

The survey area is approximately 12.566 $m^2$ per point, which we'll use as an offset in our density models.

#### Step 2: Creating Predictor Variables

We need to convert site identity into ecologically meaningful predictor variables based on site characteristics.

```
siteLevels <- factor(c("APA", "PARNA", "Secundaria"),
                      levels = c("APA", "PARNA", "Secundaria"))

# Invasive Species Presence (assuming key invasives like Cats and Tejus are largely absent from Seconda
mabuia_dd[Site == "APA", InvasiveSpeciesPresence := 1]
mabuia_dd[Site == "PARNA", InvasiveSpeciesPresence := 1]
mabuia_dd[Site == "Secundaria", InvasiveSpeciesPresence := 0]

# Food Supplementation
mabuia_dd[Site == "APA", FoodSupplementation := 1]
mabuia_dd[Site == "PARNA", FoodSupplementation := 0]
mabuia_dd[Site == "Secundaria", FoodSupplementation := 0]

# Convert to factors for modeling
mabuia_dd[, InvasiveSpeciesPresence := as.factor(InvasiveSpeciesPresence)]
mabuia_dd[, FoodSupplementation := as.factor(FoodSupplementation)]
head(mabuia_dd)
```

```
##    LocationID      Date   Time Location   Site Island          Habitat Temp_oC
##        <char>    <char> <char>   <char> <char> <char>           <char>   <int>
## 1:      IPA56 2/11/2016  16:25   3 Paus    APA   Main           Urbana      29
## 2:     IPA106 11/8/2016   8:16   Abreus  PARNA   Main   Arbustiva Alta      30
## 3:     IPA107 11/8/2016   8:30   Abreus  PARNA   Main    Arborea Baixa      30
## 4:     IPA108 11/8/2016   8:45   Abreus  PARNA   Main      Gram\xednea      30
## 5:     IPA109 11/8/2016   9:07   Abreus  PARNA   Main Pedreira (costa)      30
## 6:     IPA110 11/8/2016   9:30   Abreus  PARNA   Main Pedreira (costa)      31
##      UTM    NOR     EAS Counts ObsMin survey_area InvasiveSpeciesPresence
##    <int>  <int>   <int>  <int>  <int>       <num>                  <fctr>
## 1:    25 564037 9574144      3      7    12.56637                       1
## 2:    25 564199 9573143      1      7    12.56637                       1
## 3:    25 564452 9572968      1      7    12.56637                       1
## 4:    25 564676 9572953      1      7    12.56637                       1
## 5:    25 565228 9573056      0      7    12.56637                       1
## 6:    25 565055 9573102      0      7    12.56637                       1
##    FoodSupplementation
##                 <fctr>
## 1:                   1
## 2:                   0
## 3:                   0
## 4:                   0
## 5:                   0
## 6:                   0
```

This creates a 2x2 factorial design where we can test the independent and interactive effects of invasive species presence and food supplementation.

#### Step 3: Fitting the Main Density Model

We use a negative binomial GLM because count data often shows overdispersion (variance > mean).

```
model_hierarchical_main <- MASS::glm.nb(
  Counts ~ InvasiveSpeciesPresence + FoodSupplementation + offset(log(survey_area)),
```

```r
  data = mabuia_dd,
  na.action = na.omit
)
summary(model_hierarchical_main)
```

```
##
## Call:
## MASS::glm.nb(formula = Counts ~ InvasiveSpeciesPresence + FoodSupplementation +
##     offset(log(survey_area)), data = mabuia_dd, na.action = na.omit,
##     init.theta = 1.151932299, link = log)
##
## Coefficients:
##                          Estimate Std. Error z value Pr(>|z|)
## (Intercept)               -0.8897     0.1694  -5.252 1.51e-07 ***
## InvasiveSpeciesPresence1  -0.9666     0.2318  -4.169 3.05e-05 ***
## FoodSupplementation1       0.2687     0.2162   1.243    0.214
## ---
## Signif. codes:  0 '***' 0.001 '**' 0.01 '*' 0.05 '.' 0.1 ' ' 1
##
## (Dispersion parameter for Negative Binomial(1.1519) family taken to be 1)
##
##     Null deviance: 184.46  on 149  degrees of freedom
## Residual deviance: 165.06  on 147  degrees of freedom
## AIC: 663.54
##
## Number of Fisher Scoring iterations: 1
##
##
##               Theta:  1.152
##           Std. Err.:  0.206
##
##  2 x log-likelihood:  -655.544
```

The offset term `log(survey_area)` allows us to model density (individuals per unit area) rather than raw counts.

#### Step 4: Model Validation

Good model validation is critical for reliable inference.

```r
r2(model_hierarchical_main)
```

```
## # R2 for Generalized Linear Regression
##   Nagelkerke's R2: 0.172
```

```r
sim_res <- simulateResiduals(model_hierarchical_main)
plot(sim_res)
```

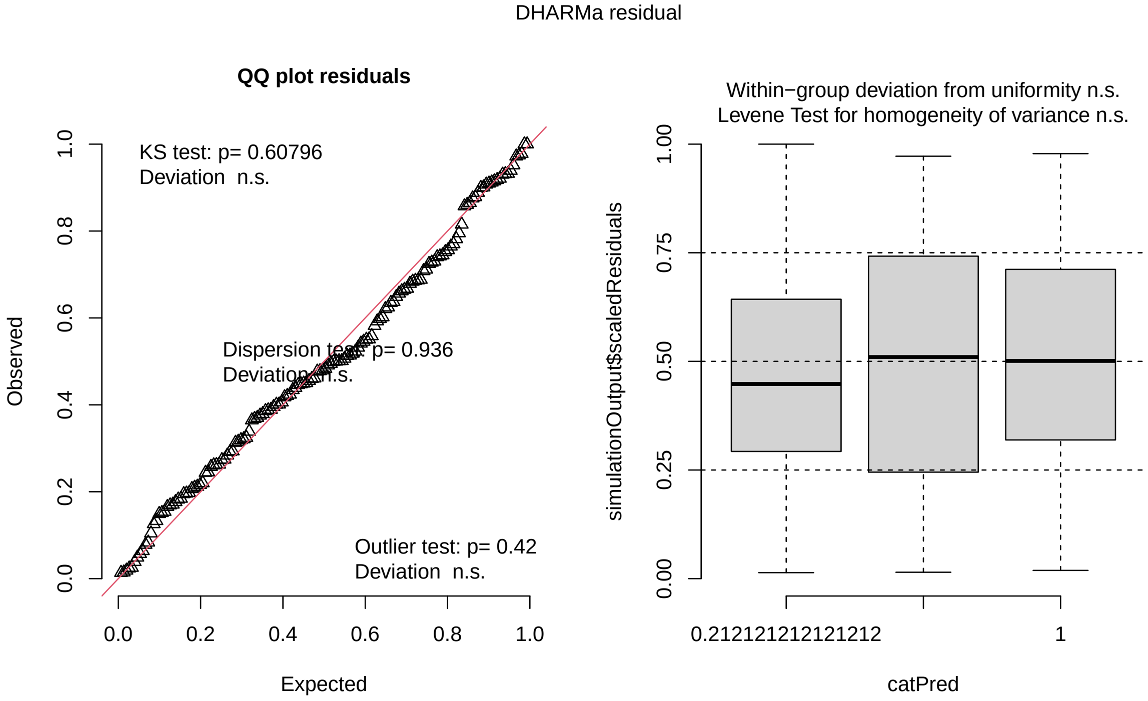

DHARMa provides robust residual diagnostics by simulating from the fitted model. Good residuals should show uniform distribution in QQ plots and no patterns in residual vs. fitted plots.

#### Step 5: Alternative Model Formulations

We test several alternative model structures to ensure we've chosen the best approach.

##### Interaction Model

```
model_interaction <- MASS::glm.nb(
  Counts ~ InvasiveSpeciesPresence * FoodSupplementation + offset(log(survey_area)),
  data = mabuia_dd
)
summary(model_interaction)
```

```
##
## Call:
## MASS::glm.nb(formula = Counts ~ InvasiveSpeciesPresence * FoodSupplementation +
##     offset(log(survey_area)), data = mabuia_dd, init.theta = 1.151932299,
##     link = log)
##
## Coefficients: (1 not defined because of singularities)
##                                               Estimate Std. Error z value
## (Intercept)                                    -0.8897     0.1694  -5.252
## InvasiveSpeciesPresence1                       -0.9666     0.2318  -4.169
## FoodSupplementation1                            0.2687     0.2162   1.243
## InvasiveSpeciesPresence1:FoodSupplementation1       NA         NA      NA
##                                               Pr(>|z|)
## (Intercept)                                   1.51e-07 ***
## InvasiveSpeciesPresence1                      3.05e-05 ***
```

```
## FoodSupplementation1                             0.214
## InvasiveSpeciesPresence1:FoodSupplementation1       NA
## ---
## Signif. codes:  0 '***' 0.001 '**' 0.01 '*' 0.05 '.' 0.1 ' ' 1
##
## (Dispersion parameter for Negative Binomial(1.1519) family taken to be 1)
##
##     Null deviance: 184.46  on 149  degrees of freedom
## Residual deviance: 165.06  on 147  degrees of freedom
## AIC: 663.54
##
## Number of Fisher Scoring iterations: 1
##
##
##               Theta:  1.152
##           Std. Err.:  0.206
##
##  2 x log-likelihood:  -655.544
```

##### Mixed Effects Models

```r
model_random <- glmmTMB::glmmTMB(
  Counts ~ InvasiveSpeciesPresence + FoodSupplementation + offset(log(survey_area)) + (1 | Site),
  family = nbinom2,
  data = mabuia_dd
)
summary(model_random)
```

```
##  Family: nbinom2  ( log )
## Formula:
## Counts ~ InvasiveSpeciesPresence + FoodSupplementation + offset(log(survey_area)) +
##     (1 | Site)
## Data: mabuia_dd
##
##      AIC      BIC   logLik deviance df.resid
##    665.5    680.6   -327.8    655.5      145
##
## Random effects:
##
## Conditional model:
##  Groups Name        Variance  Std.Dev.
##  Site   (Intercept) 7.525e-10 2.743e-05
## Number of obs: 150, groups:  Site, 3
##
## Dispersion parameter for nbinom2 family (): 1.15
##
## Conditional model:
##                          Estimate Std. Error z value Pr(>|z|)
## (Intercept)               -0.8897     0.1694  -5.252 1.51e-07 ***
## InvasiveSpeciesPresence1  -0.9666     0.2318  -4.169 3.05e-05 ***
## FoodSupplementation1       0.2687     0.2162   1.243    0.214
## ---
## Signif. codes:  0 '***' 0.001 '**' 0.01 '*' 0.05 '.' 0.1 ' ' 1
```

```r
model_random_interaction <- glmmTMB::glmmTMB(
  Counts ~ InvasiveSpeciesPresence * FoodSupplementation + offset(log(survey_area)) + (1 | Site),
  family = nbinom2,
  data = mabuia_dd
)
summary(model_random_interaction)
```

```
##  Family: nbinom2  ( log )
## Formula:
## Counts ~ InvasiveSpeciesPresence * FoodSupplementation + offset(log(survey_area)) +
##     (1 | Site)
## Data: mabuia_dd
##
##      AIC      BIC   logLik deviance df.resid
##    665.5    680.6   -327.8    655.5      145
##
## Random effects:
##
## Conditional model:
##  Groups Name        Variance  Std.Dev.
##  Site   (Intercept) 7.525e-10 2.743e-05
## Number of obs: 150, groups:  Site, 3
##
## Dispersion parameter for nbinom2 family (): 1.15
##
## Conditional model:
##                                              Estimate Std. Error z value
## (Intercept)                                   -0.8897     0.1694  -5.252
## InvasiveSpeciesPresence1                      -0.9666     0.2318  -4.169
## FoodSupplementation1                           0.2687     0.2162   1.243
## InvasiveSpeciesPresence1:FoodSupplementation1      NA         NA      NA
##                                              Pr(>|z|)
## (Intercept)                                  1.51e-07 ***
## InvasiveSpeciesPresence1                     3.05e-05 ***
## FoodSupplementation1                            0.214
## InvasiveSpeciesPresence1:FoodSupplementation1      NA
## ---
## Signif. codes:  0 '***' 0.001 '**' 0.01 '*' 0.05 '.' 0.1 ' ' 1
```

##### Model Selection

```r
model_comparison <- AIC(
  model_hierarchical_main,      # Original glm.nb model
  model_interaction,            # GLM with interaction
  model_random,                 # GLMM with random effect
  model_random_interaction      # GLMM with both interaction + random
)
print("Model AIC Comparison:")
```

```
## [1] "Model AIC Comparison:"
```

```r
print(model_comparison)
```

```
##                          df      AIC
```

```
## model_hierarchical_main   4 663.5444
## model_interaction         4 663.5444
## model_random              5 665.5444
## model_random_interaction  5 665.5444
```

**Conclusion**: The original model has the best AIC, is interpretable, answers the ecological question clearly, and passes all residual diagnostics.

#### Step 6: Estimated Marginal Means

```r
emm_density_hierarchical <- emmeans(model_hierarchical_main,
                                    specs = ~ InvasiveSpeciesPresence + FoodSupplementation,
                                    type = "response",
                                    offset = 0)

summary(emm_density_hierarchical, infer = TRUE)
```

```
##  InvasiveSpeciesPresence FoodSupplementation response     SE  df asymp.LCL
##  0                       0                      0.411 0.0696 Inf     0.295
##  1                       0                      0.156 0.0247 Inf     0.115
##  0                       1                      0.537 0.1476 Inf     0.314
##  1                       1                      0.204 0.0301 Inf     0.153
##  asymp.UCL null z.ratio p.value
##      0.573    1  -5.252  <.0001
##      0.213    1 -11.730  <.0001
##      0.921    1  -2.261  0.0238
##      0.273    1 -10.782  <.0001
##
## Confidence level used: 0.95
## Intervals are back-transformed from the log scale
## Tests are performed on the log scale
```

Setting `offset = 0` gives us density per unit area rather than total abundance.

#### Step 7: Creating the Density Visualization

##### Data Preparation for Plotting

```r
density_df <- as.data.frame(summary(emm_density_hierarchical, infer = TRUE))

# Map factor combinations to meaningful site names
density_df$Site <- ifelse(
  density_df$InvasiveSpeciesPresence == 0 & density_df$FoodSupplementation == 1, "Hypothetical Site",
  ifelse(
    density_df$InvasiveSpeciesPresence == 0 & density_df$FoodSupplementation == 0, "Secondary Islands",
    ifelse(
      density_df$InvasiveSpeciesPresence == 1 & density_df$FoodSupplementation == 1, "APA",
      "PARNAMAR"
    )
  )
)

density_df$Site <- factor(density_df$Site, levels = c("Hypothetical Site", "Secondary Islands", "APA",

# Rename columns for clarity
```

```r
names(density_df)[names(density_df) == 'response'] <- 'mean_density'
names(density_df)[names(density_df) == 'asymp.LCL'] <- 'lower_ci'
names(density_df)[names(density_df) == 'asymp.UCL'] <- 'upper_ci'
print(density_df)
```

```
##  InvasiveSpeciesPresence FoodSupplementation mean_density         SE  df
##  0                       0                      0.4107918 0.06959005 Inf
##  1                       0                      0.1562612 0.02472831 Inf
##  0                       1                      0.5374264 0.14759153 Inf
##  1                       1                      0.2044318 0.03009994 Inf
##   lower_ci  upper_ci null z.ratio p.value Site
##  0.2947300 0.5725577    1  -5.252  <.0001 Secondary Islands
##  0.1145905 0.2130854    1 -11.730  <.0001 PARNAMAR
##  0.3137303 0.9206223    1  -2.261  0.0238 Hypothetical Site
##  0.1531863 0.2728204    1 -10.782  <.0001 APA
##
## Confidence level used: 0.95
## Intervals are back-transformed from the log scale
## Tests are performed on the log scale
```

##### Statistical Significance Testing

```r
pairs_emm <- contrast(emm_density_hierarchical, method = "pairwise", adjust = "tukey")
summary_pw <- summary(pairs_emm)
p_values <- summary_pw$p.value
print(p_values)
```

```
## [1] 0.0001790051 0.5993115525 0.0101326414 0.0079318154 0.5993115525
## [6] 0.0001790051
```

##### Processing Significance Letters

```r
# Helper function to clean emmeans factor level names
clean_level_name <- function(level_string) {
  cleaned <- gsub("InvasiveSpeciesPresence", "", level_string)
  cleaned <- gsub("FoodSupplementation", "", cleaned)
  cleaned <- gsub(" ", "", cleaned)
  return(cleaned)
}

contrast_names_raw <- as.character(summary_pw$contrast)
p_value_names <- sapply(contrast_names_raw, function(contrast) {
  parts <- strsplit(contrast, " / ")[[1]]
  key1 <- clean_level_name(trimws(parts[1]))
  key2 <- clean_level_name(trimws(parts[2]))
  return(paste(key1, key2, sep = "-"))
})
names(p_values) <- p_value_names

# Generate significance letters
letters_result <- multcompLetters(p_values)
significance_letters <- letters_result$Letters

# Add letters to plotting data
```

```
density_df$key <- paste0(density_df$InvasiveSpeciesPresence, density_df$FoodSupplementation)
density_df$significance_group <- significance_letters[density_df$key]
density_df$y_position_for_text <- density_df$upper_ci + 0.05
```

##### Creating the Final Density Plot

```
two_line_labels <- c(
  "Hypothetical\n(Subsidies / Reduced Predators)",
  "Secondary Islands\n(No Subsidies / Reduced Predators)",
  "APA\n(Subsidies / Full Predators)",
  "PARNAMAR\n(No Subsidies / Full Predators)"
)

density_plot <- ggplot(density_df, aes(x = Site, y = mean_density, fill = Site)) +
  # Add bars and error bars from the GLM
  geom_bar(stat = "identity", color = "black", alpha = 0.8) +
  geom_errorbar(aes(ymin = lower_ci, ymax = upper_ci),
                width = 0.2, color = "black", linewidth = 0.5) +
  # Add significance letters from the GLM
  geom_text(aes(y = y_position_for_text, label = significance_group),
            size = 6, fontface = "bold", color = "black") +

  # --- ANNOTATION FOR CMR RESULT ---
  # Add a bracket connecting APA and PARNAMAR bars
  annotate("segment", x = "APA", xend = "PARNAMAR", y = 0.40, yend = 0.40,
           color = "black", linewidth = 0.7) +
  # New position for the left vertical tick (pointing down).
  annotate("segment", x = "APA", xend = "APA", y = 0.40, yend = 0.38,
           color = "black", linewidth = 0.7) +
  # New position for the right vertical tick (pointing down).
  annotate("segment", x = "PARNAMAR", xend = "PARNAMAR", y = 0.40, yend = 0.38,
           color = "black", linewidth = 0.7) +
  # New position for the text, slightly above the bracket line.
  annotate("text", x = 3.5, y = 0.43, label = "CMR confirms APA > PARNAMAR (p < 0.001)",
           fontface = "italic", color = "black", size = 4) +

  # --- SCALES, LABELS, and THEMING ---
  scale_x_discrete(labels = two_line_labels) +
  scale_fill_viridis_d(option = "D") +
  labs(
    y = expression("Estimated Density (individuals / m"^2*")")
  ) +
  theme_bw(base_size = 14) +
  theme(
    axis.title = element_text(face = "bold"),
    axis.text.x = element_text(face = "bold"),
    panel.grid.major.x = element_blank(),
    panel.grid.minor = element_blank(),
    legend.position = "none"
  )

print(density_plot)
```

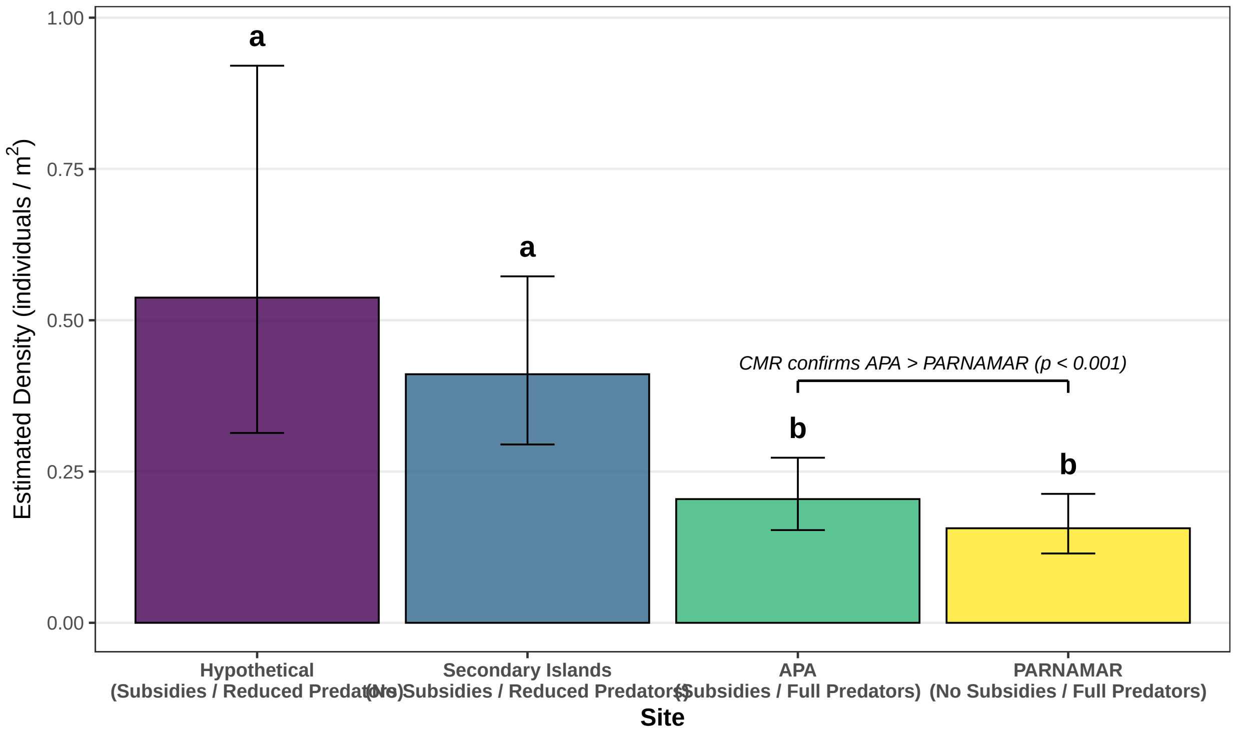

#### Mark-Recapture Analysis

Analysis was performed using the Poisson-log normal mark-resight model to separate detection probability from true abundance.

##### Step 1: Loading CMR Data

```
ch <- read.xlsx(file = file.path(getwd(), "data/CRM_Data.xlsx"),
                header = TRUE,
                sheetName = "ch",
                as.data.frame = TRUE)
ch$ch <- as.character(ch$ch)
ch$colonies <- as.factor(ch$colonies)

covars <- read.xlsx(file = file.path(getwd(), "data/CRM_Data.xlsx"),
                    header = TRUE,
                    sheetName = "covars",
                    as.data.frame = TRUE)
```

##### Step 2: Creating Model Variables

```
nocc <- 6  # Number of occasions
groupsNames <- unique(covars$colonies)
nGroups <- length(groupsNames)

# Create matrices for different count types
unmarkedSeen <- matrix(covars[, "unmarkedSeen"],
                       nrow = nGroups,
```

```
                       ncol = nocc,
                       byrow = TRUE)

markedUnidentified <- matrix(covars[, "markedUnidentified"],
                             nrow = nGroups,
                             ncol = nocc,
                             byrow = TRUE)

knownMarks <- matrix(covars[, "knownMarks"],
                     nrow = nGroups,
                     ncol = nocc,
                     byrow = TRUE)

effort <- matrix(covars[, "effort"],
                 nrow = nGroups,
                 ncol = nocc,
                 byrow = TRUE)

site <- matrix(covars[, "site"],
               nrow = nGroups,
               ncol = nocc,
               byrow = TRUE)
```

##### Step 3: Data Processing

```
mabuya.process <- process.data(ch,
                               model = "PoissonMR",
                               time.intervals = rep(1, times = (nocc-1)),
                               groups = "colonies",
                               counts = list("Unmarked Seen" = unmarkedSeen,
                                             "Marked Unidentified" = markedUnidentified,
                                             "Known Marks" = knownMarks))

mabuya.ddl <- make.design.data(mabuya.process)
```

##### Step 4: Adding Covariates

Now we include covariates such as effort and site type where they should be placed.

###### For Alpha (Detection Probability)

```
mabuya.ddl$alpha$effort <- as.numeric(apply(X = mabuya.ddl$alpha, 1, function(x){
  col <- x[["group"]]
  occ <- as.numeric(x[["time"]])
  eff <- covars[covars$colonies==col&covars$occasion==occ, "effort"]
}))

mabuya.ddl$alpha$site <- as.character(apply(X = mabuya.ddl$alpha, 1, function(x){
  col <- x[["group"]]
  occ <- as.numeric(x[["time"]])
  site <- covars[covars$colonies==col&covars$occasion==occ, "site"]
}))
```

**For U (Abundance)**

```
mabuya.ddl$U$site <- as.character(apply(X = mabuya.ddl$U, 1, function(x){
  col <- x[["group"]]
  occ <- as.numeric(x[["time"]])
  site <- covars[covars$colonies==col&covars$occasion==occ, "site"]
}))
```

#### Step 5: Defining Model Formulas

```
# Parameter formulas for alpha (detection probability)
alpha.dot <- list(formula = ~ 1)
alpha.effort <- list(formula = ~ effort)
alpha.site <- list(formula = ~ site)
alpha.additive <- list(formula = ~ effort + site)
alpha.interaction <- list(formula = ~ effort * site)

# Parameters that will be the same in all models
U.site <- list(formula = ~ site)
Phi.fixed1 <- list(formula = ~ 1, fixed = 1)
zero <- list(formula = ~ 1, fixed = 0)  # For sigma and Gammas
```

#### Step 6: Building Model List

```
model.list <- list(
  "alpha(.) U(site)" = list(
    alpha = alpha.dot,
    U = U.site,
    Phi = Phi.fixed1,
    sigma = zero,
    GammaDoublePrime = zero,
    GammaPrime = zero
  ),
  "alpha(effort) U(site)" = list(
    alpha = alpha.effort,
    U = U.site,
    Phi = Phi.fixed1,
    sigma = zero,
    GammaDoublePrime = zero,
    GammaPrime = zero
  ),
  "alpha(site) U(site)" = list(
    alpha = alpha.site,
    U = U.site,
    Phi = Phi.fixed1,
    sigma = zero,
    GammaDoublePrime = zero,
    GammaPrime = zero
  ),
  "alpha(effort+site) U(site)" = list(
    alpha = alpha.additive,
    U = U.site,
    Phi = Phi.fixed1,
```

```
    sigma = zero,
    GammaDoublePrime = zero,
    GammaPrime = zero
  ),
  "alpha(effort*site) U(site)" = list(
    alpha = alpha.interaction,
    U = U.site,
    Phi = Phi.fixed1,
    sigma = zero,
    GammaDoublePrime = zero,
    GammaPrime = zero
  )
)
```

##### Step 7: Fitting All Models

```
mabuya_model_1 <- mark(
  data = mabuya.process,
  ddl = mabuya.ddl,
  model.parameters = model.list[["alpha(.) U(site)"]],
  delete = TRUE
)
```

```
##
## Output summary for PoissonMR model
## Name : alpha(~1)sigma(~1)U(~site)Phi(~1)Gamma''(~1)Gamma'(~1)
##
## Npar :  3
## -2lnL:  872.2775
## AICc :  878.3273
##
## Beta
##                      estimate        se       lcl       ucl
## alpha:(Intercept) -1.354831 0.0940721 -1.539212 -1.170450
## U:(Intercept)      4.158624 0.1063001  3.950276  4.366973
## U:sitePARNA       -1.760433 0.1292842 -2.013830 -1.507036
##
##
## Real Parameter alpha
##                                        1         2         3         4         5
## Group:coloniesAmericano      0.2579909 0.2579909 0.2579909 0.2579909 0.2579909
## Group:coloniesCapimAcu       0.2579909 0.2579909 0.2579909 0.2579909 0.2579909
## Group:coloniesForteBoldro    0.2579909 0.2579909 0.2579909 0.2579909 0.2579909
## Group:coloniesLeao           0.2579909 0.2579909 0.2579909 0.2579909 0.2579909
## Group:coloniesPedreiraSueste 0.2579909 0.2579909 0.2579909 0.2579909 0.2579909
## Group:coloniesPiquinho       0.2579909 0.2579909 0.2579909 0.2579909 0.2579909
## Group:coloniesPraiaBoldro    0.2579909 0.2579909 0.2579909 0.2579909 0.2579909
## Group:coloniesTejuAcu        0.2579909 0.2579909 0.2579909 0.2579909 0.2579909
##                                        6
## Group:coloniesAmericano      0.2579909
## Group:coloniesCapimAcu       0.2579909
## Group:coloniesForteBoldro    0.2579909
## Group:coloniesLeao           0.2579909
```

```
## Group:coloniesPedreiraSueste 0.2579909
## Group:coloniesPiquinho       0.2579909
## Group:coloniesPraiaBoldro    0.2579909
## Group:coloniesTejuAcu        0.2579909
##
##
## Real Parameter sigma
##                               1  2  3  4  5  6
## Group:coloniesAmericano      NA NA NA NA NA NA
## Group:coloniesCapimAcu       NA NA NA NA NA NA
## Group:coloniesForteBoldro    NA NA NA NA NA NA
## Group:coloniesLeao           NA NA NA NA NA NA
## Group:coloniesPedreiraSueste NA NA NA NA NA NA
## Group:coloniesPiquinho       NA NA NA NA NA NA
## Group:coloniesPraiaBoldro    NA NA NA NA NA NA
## Group:coloniesTejuAcu        NA NA NA NA NA NA
##
##
## Real Parameter U
##                                     1        2        3        4        5
## Group:coloniesAmericano      63.98345 63.98345 63.98345 63.98345 63.98345
## Group:coloniesCapimAcu       11.00326 11.00326 11.00326 11.00326 11.00326
## Group:coloniesForteBoldro    63.98345 63.98345 63.98345 63.98345 63.98345
## Group:coloniesLeao           11.00326 11.00326 11.00326 11.00326 11.00326
## Group:coloniesPedreiraSueste 11.00326 11.00326 11.00326 11.00326 11.00326
## Group:coloniesPiquinho       11.00326 11.00326 11.00326 11.00326 11.00326
## Group:coloniesPraiaBoldro    63.98345 63.98345 63.98345 63.98345 63.98345
## Group:coloniesTejuAcu        63.98345 63.98345 63.98345 63.98345 63.98345
##                                     6
## Group:coloniesAmericano      63.98345
## Group:coloniesCapimAcu       11.00326
## Group:coloniesForteBoldro    63.98345
## Group:coloniesLeao           11.00326
## Group:coloniesPedreiraSueste 11.00326
## Group:coloniesPiquinho       11.00326
## Group:coloniesPraiaBoldro    63.98345
## Group:coloniesTejuAcu        63.98345
##
##
## Real Parameter Phi
## Group:coloniesAmericano
##   1 2 3 4 5
## 1
## 2
## 3
## 4
## 5
##
## Group:coloniesCapimAcu
##   1 2 3 4 5
## 1
## 2
## 3
## 4
```

```
## 5
##
## Group:coloniesForteBoldro
##   1 2 3 4 5
## 1
## 2
## 3
## 4
## 5
##
## Group:coloniesLeao
##   1 2 3 4 5
## 1
## 2
## 3
## 4
## 5
##
## Group:coloniesPedreiraSueste
##   1 2 3 4 5
## 1
## 2
## 3
## 4
## 5
##
## Group:coloniesPiquinho
##   1 2 3 4 5
## 1
## 2
## 3
## 4
## 5
##
## Group:coloniesPraiaBoldro
##   1 2 3 4 5
## 1
## 2
## 3
## 4
## 5
##
## Group:coloniesTejuAcu
##   1 2 3 4 5
## 1
## 2
## 3
## 4
## 5
##
##
## Real Parameter GammaDoublePrime
## Group:coloniesAmericano
##   1 2 3 4 5
```

```
## 1
## 2
## 3
## 4
## 5
##
## Group:coloniesCapimAcu
##   1 2 3 4 5
## 1
## 2
## 3
## 4
## 5
##
## Group:coloniesForteBoldro
##   1 2 3 4 5
## 1
## 2
## 3
## 4
## 5
##
## Group:coloniesLeao
##   1 2 3 4 5
## 1
## 2
## 3
## 4
## 5
##
## Group:coloniesPedreiraSueste
##   1 2 3 4 5
## 1
## 2
## 3
## 4
## 5
##
## Group:coloniesPiquinho
##   1 2 3 4 5
## 1
## 2
## 3
## 4
## 5
##
## Group:coloniesPraiaBoldro
##   1 2 3 4 5
## 1
## 2
## 3
## 4
## 5
##
```

```
## Group:coloniesTejuAcu
##   1 2 3 4 5
## 1
## 2
## 3
## 4
## 5
##
##
## Real Parameter GammaPrime
## Group:coloniesAmericano
##   2 3 4 5
## 1
## 2
## 3
## 4
##
## Group:coloniesCapimAcu
##   2 3 4 5
## 1
## 2
## 3
## 4
##
## Group:coloniesForteBoldro
##   2 3 4 5
## 1
## 2
## 3
## 4
##
## Group:coloniesLeao
##   2 3 4 5
## 1
## 2
## 3
## 4
##
## Group:coloniesPedreiraSueste
##   2 3 4 5
## 1
## 2
## 3
## 4
##
## Group:coloniesPiquinho
##   2 3 4 5
## 1
## 2
## 3
## 4
##
## Group:coloniesPraiaBoldro
##   2 3 4 5
```

```
## 1
## 2
## 3
## 4
##
## Group:coloniesTejuAcu
##   2 3 4 5
## 1
## 2
## 3
## 4
```

```r
mabuya_model_2 <- mark(
  data = mabuya.process,
  ddl = mabuya.ddl,
  model.parameters = model.list[["alpha(effort) U(site)"]],
  delete = TRUE
)
```

```
##
## Output summary for PoissonMR model
## Name : alpha(~effort)sigma(~1)U(~site)Phi(~1)Gamma''(~1)Gamma'(~1)
##
## Npar :  4
## -2lnL:  849.1093
## AICc :  857.1925
##
## Beta
##                       estimate        se        lcl        ucl
## alpha:(Intercept) -1.8070139 0.1359359 -2.0734483 -1.5405795
## alpha:effort       0.0189359 0.0039292  0.0112346  0.0266371
## U:(Intercept)      4.1289170 0.1067975  3.9195938  4.3382401
## U:sitePARNA       -1.5993993 0.1343945 -1.8628125 -1.3359862
##
##
## Real Parameter alpha
##                                        1         2         3         4         5
## Group:coloniesAmericano      0.4230688 0.2180620 0.2635223 0.2397169 0.2180620
## Group:coloniesCapimAcu       0.2442994 0.2021552 0.2180620 0.2180620 0.2180620
## Group:coloniesForteBoldro    0.3184600 0.2180620 0.2635223 0.2635223 0.3008733
## Group:coloniesLeao           0.2308082 0.2397169 0.2099580 0.2264788 0.2099580
## Group:coloniesPedreiraSueste 0.2397169 0.2264788 0.2442994 0.2308082 0.2222305
## Group:coloniesPiquinho       0.2308082 0.2060197 0.2736938 0.2180620 0.2180620
## Group:coloniesPraiaBoldro    0.2308082 0.3008733 0.3500851 0.2789258 0.2060197
## Group:coloniesTejuAcu        0.2180620 0.1909914 0.1909914 0.1983633 0.1983633
##                                        6
## Group:coloniesAmericano      0.2180620
## Group:coloniesCapimAcu       0.2352204
## Group:coloniesForteBoldro    0.2635223
## Group:coloniesLeao           0.2180620
## Group:coloniesPedreiraSueste 0.2139716
## Group:coloniesPiquinho       0.1983633
## Group:coloniesPraiaBoldro    0.3500851
## Group:coloniesTejuAcu        0.1983633
##
```

```
##
## Real Parameter sigma
##                               1  2  3  4  5  6
## Group:coloniesAmericano      NA NA NA NA NA NA
## Group:coloniesCapimAcu       NA NA NA NA NA NA
## Group:coloniesForteBoldro    NA NA NA NA NA NA
## Group:coloniesLeao           NA NA NA NA NA NA
## Group:coloniesPedreiraSueste NA NA NA NA NA NA
## Group:coloniesPiquinho       NA NA NA NA NA NA
## Group:coloniesPraiaBoldro    NA NA NA NA NA NA
## Group:coloniesTejuAcu        NA NA NA NA NA NA
##
##
## Real Parameter U
##                                     1        2        3        4        5
## Group:coloniesAmericano      62.11062 62.11062 62.11062 62.11062 62.11062
## Group:coloniesCapimAcu       12.54745 12.54745 12.54745 12.54745 12.54745
## Group:coloniesForteBoldro    62.11062 62.11062 62.11062 62.11062 62.11062
## Group:coloniesLeao           12.54745 12.54745 12.54745 12.54745 12.54745
## Group:coloniesPedreiraSueste 12.54745 12.54745 12.54745 12.54745 12.54745
## Group:coloniesPiquinho       12.54745 12.54745 12.54745 12.54745 12.54745
## Group:coloniesPraiaBoldro    62.11062 62.11062 62.11062 62.11062 62.11062
## Group:coloniesTejuAcu        62.11062 62.11062 62.11062 62.11062 62.11062
##                                     6
## Group:coloniesAmericano      62.11062
## Group:coloniesCapimAcu       12.54745
## Group:coloniesForteBoldro    62.11062
## Group:coloniesLeao           12.54745
## Group:coloniesPedreiraSueste 12.54745
## Group:coloniesPiquinho       12.54745
## Group:coloniesPraiaBoldro    62.11062
## Group:coloniesTejuAcu        62.11062
##
##
## Real Parameter Phi
## Group:coloniesAmericano
##   1 2 3 4 5
## 1
## 2
## 3
## 4
## 5
##
## Group:coloniesCapimAcu
##   1 2 3 4 5
## 1
## 2
## 3
## 4
## 5
##
## Group:coloniesForteBoldro
##   1 2 3 4 5
## 1
```

```
## 2
## 3
## 4
## 5
##
## Group:coloniesLeao
##   1 2 3 4 5
## 1
## 2
## 3
## 4
## 5
##
## Group:coloniesPedreiraSueste
##   1 2 3 4 5
## 1
## 2
## 3
## 4
## 5
##
## Group:coloniesPiquinho
##   1 2 3 4 5
## 1
## 2
## 3
## 4
## 5
##
## Group:coloniesPraiaBoldro
##   1 2 3 4 5
## 1
## 2
## 3
## 4
## 5
##
## Group:coloniesTejuAcu
##   1 2 3 4 5
## 1
## 2
## 3
## 4
## 5
##
##
## Real Parameter GammaDoublePrime
## Group:coloniesAmericano
##   1 2 3 4 5
## 1
## 2
## 3
## 4
## 5
```

```
##
## Group:coloniesCapimAcu
##   1 2 3 4 5
## 1
## 2
## 3
## 4
## 5
##
## Group:coloniesForteBoldro
##   1 2 3 4 5
## 1
## 2
## 3
## 4
## 5
##
## Group:coloniesLeao
##   1 2 3 4 5
## 1
## 2
## 3
## 4
## 5
##
## Group:coloniesPedreiraSueste
##   1 2 3 4 5
## 1
## 2
## 3
## 4
## 5
##
## Group:coloniesPiquinho
##   1 2 3 4 5
## 1
## 2
## 3
## 4
## 5
##
## Group:coloniesPraiaBoldro
##   1 2 3 4 5
## 1
## 2
## 3
## 4
## 5
##
## Group:coloniesTejuAcu
##   1 2 3 4 5
## 1
## 2
## 3
```

```
## 4
## 5
##
##
## Real Parameter GammaPrime
## Group:coloniesAmericano
##   2 3 4 5
## 1
## 2
## 3
## 4
##
## Group:coloniesCapimAcu
##   2 3 4 5
## 1
## 2
## 3
## 4
##
## Group:coloniesForteBoldro
##   2 3 4 5
## 1
## 2
## 3
## 4
##
## Group:coloniesLeao
##   2 3 4 5
## 1
## 2
## 3
## 4
##
## Group:coloniesPedreiraSueste
##   2 3 4 5
## 1
## 2
## 3
## 4
##
## Group:coloniesPiquinho
##   2 3 4 5
## 1
## 2
## 3
## 4
##
## Group:coloniesPraiaBoldro
##   2 3 4 5
## 1
## 2
## 3
## 4
##
```

```
## Group:coloniesTejuAcu
##   2 3 4 5
## 1
## 2
## 3
## 4
```

```
mabuya_model_3 <- mark(
  data = mabuya.process,
  ddl = mabuya.ddl,
  model.parameters = model.list[["alpha(site) U(site)"]],
  delete = TRUE
)
```

```
##
## Output summary for PoissonMR model
## Name : alpha(~site)sigma(~1)U(~site)Phi(~1)Gamma''(~1)Gamma'(~1)
##
## Npar :  4
## -2lnL:  871.1816
## AICc :  879.2648
##
## Beta
##                       estimate        se        lcl        ucl
## alpha:(Intercept) -1.4182458 0.1139607 -1.6416087 -1.1948830
## alpha:sitePARNA    0.2142728 0.2019031 -0.1814573  0.6100029
## U:(Intercept)      4.2220392 0.1242465  3.9785161  4.4655623
## U:sitePARNA       -1.9747053 0.2397484 -2.4446121 -1.5047985
##
##
## Real Parameter alpha
##                                        1         2         3         4         5
## Group:coloniesAmericano      0.2421384 0.2421384 0.2421384 0.2421384 0.2421384
## Group:coloniesCapimAcu       0.2999999 0.2999999 0.2999999 0.2999999 0.2999999
## Group:coloniesForteBoldro    0.2421384 0.2421384 0.2421384 0.2421384 0.2421384
## Group:coloniesLeao           0.2999999 0.2999999 0.2999999 0.2999999 0.2999999
## Group:coloniesPedreiraSueste 0.2999999 0.2999999 0.2999999 0.2999999 0.2999999
## Group:coloniesPiquinho       0.2999999 0.2999999 0.2999999 0.2999999 0.2999999
## Group:coloniesPraiaBoldro    0.2421384 0.2421384 0.2421384 0.2421384 0.2421384
## Group:coloniesTejuAcu        0.2421384 0.2421384 0.2421384 0.2421384 0.2421384
##                                        6
## Group:coloniesAmericano      0.2421384
## Group:coloniesCapimAcu       0.2999999
## Group:coloniesForteBoldro    0.2421384
## Group:coloniesLeao           0.2999999
## Group:coloniesPedreiraSueste 0.2999999
## Group:coloniesPiquinho       0.2999999
## Group:coloniesPraiaBoldro    0.2421384
## Group:coloniesTejuAcu        0.2421384
##
##
## Real Parameter sigma
##                               1  2  3  4  5  6
## Group:coloniesAmericano      NA NA NA NA NA NA
## Group:coloniesCapimAcu       NA NA NA NA NA NA
```

```
## Group:coloniesForteBoldro    NA NA NA NA NA NA
## Group:coloniesLeao           NA NA NA NA NA NA
## Group:coloniesPedreiraSueste NA NA NA NA NA NA
## Group:coloniesPiquinho       NA NA NA NA NA NA
## Group:coloniesPraiaBoldro    NA NA NA NA NA NA
## Group:coloniesTejuAcu        NA NA NA NA NA NA
##
##
## Real Parameter U
##                                      1         2         3         4         5
## Group:coloniesAmericano      68.172360 68.172360 68.172360 68.172360 68.172360
## Group:coloniesCapimAcu        9.462474  9.462474  9.462474  9.462474  9.462474
## Group:coloniesForteBoldro    68.172360 68.172360 68.172360 68.172360 68.172360
## Group:coloniesLeao            9.462474  9.462474  9.462474  9.462474  9.462474
## Group:coloniesPedreiraSueste  9.462474  9.462474  9.462474  9.462474  9.462474
## Group:coloniesPiquinho        9.462474  9.462474  9.462474  9.462474  9.462474
## Group:coloniesPraiaBoldro    68.172360 68.172360 68.172360 68.172360 68.172360
## Group:coloniesTejuAcu        68.172360 68.172360 68.172360 68.172360 68.172360
##                                      6
## Group:coloniesAmericano      68.172360
## Group:coloniesCapimAcu        9.462474
## Group:coloniesForteBoldro    68.172360
## Group:coloniesLeao            9.462474
## Group:coloniesPedreiraSueste  9.462474
## Group:coloniesPiquinho        9.462474
## Group:coloniesPraiaBoldro    68.172360
## Group:coloniesTejuAcu        68.172360
##
##
## Real Parameter Phi
## Group:coloniesAmericano
##   1 2 3 4 5
## 1
## 2
## 3
## 4
## 5
##
## Group:coloniesCapimAcu
##   1 2 3 4 5
## 1
## 2
## 3
## 4
## 5
##
## Group:coloniesForteBoldro
##   1 2 3 4 5
## 1
## 2
## 3
## 4
## 5
##
```

```
## Group:coloniesLeao
##   1 2 3 4 5
## 1
## 2
## 3
## 4
## 5
##
## Group:coloniesPedreiraSueste
##   1 2 3 4 5
## 1
## 2
## 3
## 4
## 5
##
## Group:coloniesPiquinho
##   1 2 3 4 5
## 1
## 2
## 3
## 4
## 5
##
## Group:coloniesPraiaBoldro
##   1 2 3 4 5
## 1
## 2
## 3
## 4
## 5
##
## Group:coloniesTejuAcu
##   1 2 3 4 5
## 1
## 2
## 3
## 4
## 5
##
##
## Real Parameter GammaDoublePrime
## Group:coloniesAmericano
##   1 2 3 4 5
## 1
## 2
## 3
## 4
## 5
##
## Group:coloniesCapimAcu
##   1 2 3 4 5
## 1
## 2
```

```
## 3
## 4
## 5
##
## Group:coloniesForteBoldro
##   1 2 3 4 5
## 1
## 2
## 3
## 4
## 5
##
## Group:coloniesLeao
##   1 2 3 4 5
## 1
## 2
## 3
## 4
## 5
##
## Group:coloniesPedreiraSueste
##   1 2 3 4 5
## 1
## 2
## 3
## 4
## 5
##
## Group:coloniesPiquinho
##   1 2 3 4 5
## 1
## 2
## 3
## 4
## 5
##
## Group:coloniesPraiaBoldro
##   1 2 3 4 5
## 1
## 2
## 3
## 4
## 5
##
## Group:coloniesTejuAcu
##   1 2 3 4 5
## 1
## 2
## 3
## 4
## 5
##
##
## Real Parameter GammaPrime
```

```
## Group:coloniesAmericano
##   2 3 4 5
## 1
## 2
## 3
## 4
##
## Group:coloniesCapimAcu
##   2 3 4 5
## 1
## 2
## 3
## 4
##
## Group:coloniesForteBoldro
##   2 3 4 5
## 1
## 2
## 3
## 4
##
## Group:coloniesLeao
##   2 3 4 5
## 1
## 2
## 3
## 4
##
## Group:coloniesPedreiraSueste
##   2 3 4 5
## 1
## 2
## 3
## 4
##
## Group:coloniesPiquinho
##   2 3 4 5
## 1
## 2
## 3
## 4
##
## Group:coloniesPraiaBoldro
##   2 3 4 5
## 1
## 2
## 3
## 4
##
## Group:coloniesTejuAcu
##   2 3 4 5
## 1
## 2
## 3
```

```
## 4
```

```r
mabuya_model_4 <- mark(
  data = mabuya.process,
  ddl = mabuya.ddl,
  model.parameters = model.list[["alpha(effort+site) U(site)"]],
  delete = TRUE
)
```

```
##
## Output summary for PoissonMR model
## Name : alpha(~effort + site)sigma(~1)U(~site)Phi(~1)Gamma''(~1)Gamma'(~1)
##
## Npar :  5
## -2lnL:  845.2658
## AICc :  855.3908
##
## Beta
##                       estimate        se        lcl        ucl
## alpha:(Intercept) -1.9612543 0.1597802 -2.2744234 -1.6480852
## alpha:effort       0.0205160 0.0040357  0.0126060  0.0284260
## alpha:sitePARNA    0.4154279 0.2066882  0.0103191  0.8205367
## U:(Intercept)      4.2427291 0.1249664  3.9977950  4.4876632
## U:sitePARNA       -2.0009952 0.2402727 -2.4719298 -1.5300606
##
##
## Real Parameter alpha
##                                       1         2         3         4         5
## Group:coloniesAmericano      0.3924079 0.1913762 0.2349567 0.2120498 0.1913762
## Group:coloniesCapimAcu       0.3279186 0.2670952 0.2899386 0.2899386 0.2899386
## Group:coloniesForteBoldro    0.2884615 0.1913762 0.2349567 0.2349567 0.2712426
## Group:coloniesLeao           0.3083444 0.3212595 0.2782826 0.3020828 0.2782826
## Group:coloniesPedreiraSueste 0.3212595 0.3020828 0.3279186 0.3083444 0.2959484
## Group:coloniesPiquinho       0.3083444 0.2726315 0.3708736 0.2899386 0.2899386
## Group:coloniesPraiaBoldro    0.2035251 0.2712426 0.3196229 0.2498722 0.1799525
## Group:coloniesTejuAcu        0.1913762 0.1657746 0.1657746 0.1727182 0.1727182
##                                       6
## Group:coloniesAmericano      0.1913762
## Group:coloniesCapimAcu       0.3147357
## Group:coloniesForteBoldro    0.2349567
## Group:coloniesLeao           0.2899386
## Group:coloniesPedreiraSueste 0.2840508
## Group:coloniesPiquinho       0.2616713
## Group:coloniesPraiaBoldro    0.3196229
## Group:coloniesTejuAcu        0.1727182
##
##
## Real Parameter sigma
##                               1  2  3  4  5  6
## Group:coloniesAmericano      NA NA NA NA NA NA
## Group:coloniesCapimAcu       NA NA NA NA NA NA
## Group:coloniesForteBoldro    NA NA NA NA NA NA
## Group:coloniesLeao           NA NA NA NA NA NA
## Group:coloniesPedreiraSueste NA NA NA NA NA NA
## Group:coloniesPiquinho       NA NA NA NA NA NA
```

```
## Group:coloniesPraiaBoldro    NA NA NA NA NA NA
## Group:coloniesTejuAcu        NA NA NA NA NA NA
##
##
## Real Parameter U
##                                      1         2         3         4         5
## Group:coloniesAmericano      69.597532 69.597532 69.597532 69.597532 69.597532
## Group:coloniesCapimAcu        9.409632  9.409632  9.409632  9.409632  9.409632
## Group:coloniesForteBoldro    69.597532 69.597532 69.597532 69.597532 69.597532
## Group:coloniesLeao            9.409632  9.409632  9.409632  9.409632  9.409632
## Group:coloniesPedreiraSueste  9.409632  9.409632  9.409632  9.409632  9.409632
## Group:coloniesPiquinho        9.409632  9.409632  9.409632  9.409632  9.409632
## Group:coloniesPraiaBoldro    69.597532 69.597532 69.597532 69.597532 69.597532
## Group:coloniesTejuAcu        69.597532 69.597532 69.597532 69.597532 69.597532
##                                      6
## Group:coloniesAmericano      69.597532
## Group:coloniesCapimAcu        9.409632
## Group:coloniesForteBoldro    69.597532
## Group:coloniesLeao            9.409632
## Group:coloniesPedreiraSueste  9.409632
## Group:coloniesPiquinho        9.409632
## Group:coloniesPraiaBoldro    69.597532
## Group:coloniesTejuAcu        69.597532
##
##
## Real Parameter Phi
## Group:coloniesAmericano
##   1 2 3 4 5
## 1
## 2
## 3
## 4
## 5
##
## Group:coloniesCapimAcu
##   1 2 3 4 5
## 1
## 2
## 3
## 4
## 5
##
## Group:coloniesForteBoldro
##   1 2 3 4 5
## 1
## 2
## 3
## 4
## 5
##
## Group:coloniesLeao
##   1 2 3 4 5
## 1
## 2
```

```
## 3
## 4
## 5
##
## Group:coloniesPedreiraSueste
##   1 2 3 4 5
## 1
## 2
## 3
## 4
## 5
##
## Group:coloniesPiquinho
##   1 2 3 4 5
## 1
## 2
## 3
## 4
## 5
##
## Group:coloniesPraiaBoldro
##   1 2 3 4 5
## 1
## 2
## 3
## 4
## 5
##
## Group:coloniesTejuAcu
##   1 2 3 4 5
## 1
## 2
## 3
## 4
## 5
##
##
## Real Parameter GammaDoublePrime
## Group:coloniesAmericano
##   1 2 3 4 5
## 1
## 2
## 3
## 4
## 5
##
## Group:coloniesCapimAcu
##   1 2 3 4 5
## 1
## 2
## 3
## 4
## 5
##
```

```
## Group:coloniesForteBoldro
##   1 2 3 4 5
## 1
## 2
## 3
## 4
## 5
##
## Group:coloniesLeao
##   1 2 3 4 5
## 1
## 2
## 3
## 4
## 5
##
## Group:coloniesPedreiraSueste
##   1 2 3 4 5
## 1
## 2
## 3
## 4
## 5
##
## Group:coloniesPiquinho
##   1 2 3 4 5
## 1
## 2
## 3
## 4
## 5
##
## Group:coloniesPraiaBoldro
##   1 2 3 4 5
## 1
## 2
## 3
## 4
## 5
##
## Group:coloniesTejuAcu
##   1 2 3 4 5
## 1
## 2
## 3
## 4
## 5
##
##
## Real Parameter GammaPrime
## Group:coloniesAmericano
##   2 3 4 5
## 1
## 2
```

```
## 3
## 4
##
## Group:coloniesCapimAcu
##   2 3 4 5
## 1
## 2
## 3
## 4
##
## Group:coloniesForteBoldro
##   2 3 4 5
## 1
## 2
## 3
## 4
##
## Group:coloniesLeao
##   2 3 4 5
## 1
## 2
## 3
## 4
##
## Group:coloniesPedreiraSueste
##   2 3 4 5
## 1
## 2
## 3
## 4
##
## Group:coloniesPiquinho
##   2 3 4 5
## 1
## 2
## 3
## 4
##
## Group:coloniesPraiaBoldro
##   2 3 4 5
## 1
## 2
## 3
## 4
##
## Group:coloniesTejuAcu
##   2 3 4 5
## 1
## 2
## 3
## 4
```

```
mabuya_model_5 <- mark(
  data = mabuya.process,
```

```
  ddl = mabuya.ddl,
  model.parameters = model.list[["alpha(effort*site) U(site)"]],
  delete = TRUE
)
```

```
##
## Output summary for PoissonMR model
## Name : alpha(~effort * site)sigma(~1)U(~site)Phi(~1)Gamma''(~1)Gamma'(~1)
##
## Npar :  6
## -2lnL:  845.0471
## AICc :  857.2225
##
## Beta
##                          estimate        se        lcl        ucl
## alpha:(Intercept)      -1.9710750 0.1612669 -2.2871581 -1.6549919
## alpha:effort            0.0208696 0.0041051  0.0128237  0.0289156
## alpha:sitePARNA         0.5931867 0.4306514 -0.2508901  1.4372636
## alpha:effort:sitePARNA -0.0103921 0.0221580 -0.0538218  0.0330375
## U:(Intercept)           4.2434972 0.1249911  3.9985146  4.4884797
## U:sitePARNA            -2.0002849 0.2403150 -2.4713023 -1.5292675
##
##
## Real Parameter alpha
##                                        1         2         3         4         5
## Group:coloniesAmericano      0.3955042 0.1905138 0.2347265 0.2114678 0.1905138
## Group:coloniesCapimAcu       0.3141575 0.2829074 0.2950160 0.2950160 0.2950160
## Group:coloniesForteBoldro    0.2891996 0.1905138 0.2347265 0.2347265 0.2716483
## Group:coloniesLeao           0.3044364 0.3108831 0.2888983 0.3012633 0.2888983
## Group:coloniesPedreiraSueste 0.3108831 0.3012633 0.3141575 0.3044364 0.2981233
## Group:coloniesPiquinho       0.3044364 0.2858871 0.3345411 0.2950160 0.2950160
## Group:coloniesPraiaBoldro    0.2028230 0.2716483 0.3210077 0.2498922 0.1789517
## Group:coloniesTejuAcu        0.1905138 0.1646196 0.1646196 0.1716361 0.1716361
##                                        6
## Group:coloniesAmericano      0.1905138
## Group:coloniesCapimAcu       0.3076429
## Group:coloniesForteBoldro    0.2347265
## Group:coloniesLeao           0.2950160
## Group:coloniesPedreiraSueste 0.2919411
## Group:coloniesPiquinho       0.2799587
## Group:coloniesPraiaBoldro    0.3210077
## Group:coloniesTejuAcu        0.1716361
##
##
## Real Parameter sigma
##                               1  2  3  4  5  6
## Group:coloniesAmericano      NA NA NA NA NA NA
## Group:coloniesCapimAcu       NA NA NA NA NA NA
## Group:coloniesForteBoldro    NA NA NA NA NA NA
## Group:coloniesLeao           NA NA NA NA NA NA
## Group:coloniesPedreiraSueste NA NA NA NA NA NA
## Group:coloniesPiquinho       NA NA NA NA NA NA
## Group:coloniesPraiaBoldro    NA NA NA NA NA NA
## Group:coloniesTejuAcu        NA NA NA NA NA NA
```

```
##
##
## Real Parameter U
##                                   1        2        3        4        5
## Group:coloniesAmericano      69.651008 69.651008 69.651008 69.651008 69.651008
## Group:coloniesCapimAcu        9.423554  9.423554  9.423554  9.423554  9.423554
## Group:coloniesForteBoldro    69.651008 69.651008 69.651008 69.651008 69.651008
## Group:coloniesLeao            9.423554  9.423554  9.423554  9.423554  9.423554
## Group:coloniesPedreiraSueste  9.423554  9.423554  9.423554  9.423554  9.423554
## Group:coloniesPiquinho        9.423554  9.423554  9.423554  9.423554  9.423554
## Group:coloniesPraiaBoldro    69.651008 69.651008 69.651008 69.651008 69.651008
## Group:coloniesTejuAcu        69.651008 69.651008 69.651008 69.651008 69.651008
##                                   6
## Group:coloniesAmericano      69.651008
## Group:coloniesCapimAcu        9.423554
## Group:coloniesForteBoldro    69.651008
## Group:coloniesLeao            9.423554
## Group:coloniesPedreiraSueste  9.423554
## Group:coloniesPiquinho        9.423554
## Group:coloniesPraiaBoldro    69.651008
## Group:coloniesTejuAcu        69.651008
##
##
## Real Parameter Phi
## Group:coloniesAmericano
##   1 2 3 4 5
## 1
## 2
## 3
## 4
## 5
##
## Group:coloniesCapimAcu
##   1 2 3 4 5
## 1
## 2
## 3
## 4
## 5
##
## Group:coloniesForteBoldro
##   1 2 3 4 5
## 1
## 2
## 3
## 4
## 5
##
## Group:coloniesLeao
##   1 2 3 4 5
## 1
## 2
## 3
## 4
```

```
## 5
##
## Group:coloniesPedreiraSueste
##   1 2 3 4 5
## 1
## 2
## 3
## 4
## 5
##
## Group:coloniesPiquinho
##   1 2 3 4 5
## 1
## 2
## 3
## 4
## 5
##
## Group:coloniesPraiaBoldro
##   1 2 3 4 5
## 1
## 2
## 3
## 4
## 5
##
## Group:coloniesTejuAcu
##   1 2 3 4 5
## 1
## 2
## 3
## 4
## 5
##
##
## Real Parameter GammaDoublePrime
## Group:coloniesAmericano
##   1 2 3 4 5
## 1
## 2
## 3
## 4
## 5
##
## Group:coloniesCapimAcu
##   1 2 3 4 5
## 1
## 2
## 3
## 4
## 5
##
## Group:coloniesForteBoldro
##   1 2 3 4 5
```

```
## 1
## 2
## 3
## 4
## 5
##
## Group:coloniesLeao
##   1 2 3 4 5
## 1
## 2
## 3
## 4
## 5
##
## Group:coloniesPedreiraSueste
##   1 2 3 4 5
## 1
## 2
## 3
## 4
## 5
##
## Group:coloniesPiquinho
##   1 2 3 4 5
## 1
## 2
## 3
## 4
## 5
##
## Group:coloniesPraiaBoldro
##   1 2 3 4 5
## 1
## 2
## 3
## 4
## 5
##
## Group:coloniesTejuAcu
##   1 2 3 4 5
## 1
## 2
## 3
## 4
## 5
##
##
## Real Parameter GammaPrime
## Group:coloniesAmericano
##   2 3 4 5
## 1
## 2
## 3
## 4
```

```
##
## Group:coloniesCapimAcu
##   2 3 4 5
## 1
## 2
## 3
## 4
##
## Group:coloniesForteBoldro
##   2 3 4 5
## 1
## 2
## 3
## 4
##
## Group:coloniesLeao
##   2 3 4 5
## 1
## 2
## 3
## 4
##
## Group:coloniesPedreiraSueste
##   2 3 4 5
## 1
## 2
## 3
## 4
##
## Group:coloniesPiquinho
##   2 3 4 5
## 1
## 2
## 3
## 4
##
## Group:coloniesPraiaBoldro
##   2 3 4 5
## 1
## 2
## 3
## 4
##
## Group:coloniesTejuAcu
##   2 3 4 5
## 1
## 2
## 3
## 4
```

##### Step 8: Model Selection and Best Model Results

```r
tb <- collect.models(lx = c("mabuya_model_1", "mabuya_model_2",
                            "mabuya_model_3", "mabuya_model_4",
                            "mabuya_model_5"))

tbDT <- data.table(print(tb))
```

```
##                                                                 model npar
## 4 alpha(~effort + site)sigma(~1)U(~site)Phi(~1)Gamma''(~1)Gamma'(~1)    5
## 2        alpha(~effort)sigma(~1)U(~site)Phi(~1)Gamma''(~1)Gamma'(~1)    4
## 5 alpha(~effort * site)sigma(~1)U(~site)Phi(~1)Gamma''(~1)Gamma'(~1)    6
## 1             alpha(~1)sigma(~1)U(~site)Phi(~1)Gamma''(~1)Gamma'(~1)    3
## 3          alpha(~site)sigma(~1)U(~site)Phi(~1)Gamma''(~1)Gamma'(~1)    4
##       AICc DeltaAICc       weight Deviance
## 4 855.3908  0.000000 5.535834e-01 845.2658
## 2 857.1925  1.801710 2.248779e-01 849.1093
## 5 857.2225  1.831715 2.215293e-01 845.0471
## 1 878.3273 22.936543 5.788635e-06 872.2775
## 3 879.2648 23.874000 3.622512e-06 871.1816
```

```r
print("Model selection table:")
```

```
## [1] "Model selection table:"
```

```r
tbDT[, modelPosition := rownames(print(tb))]
```

```
##                                                                 model npar
## 4 alpha(~effort + site)sigma(~1)U(~site)Phi(~1)Gamma''(~1)Gamma'(~1)    5
## 2        alpha(~effort)sigma(~1)U(~site)Phi(~1)Gamma''(~1)Gamma'(~1)    4
## 5 alpha(~effort * site)sigma(~1)U(~site)Phi(~1)Gamma''(~1)Gamma'(~1)    6
## 1             alpha(~1)sigma(~1)U(~site)Phi(~1)Gamma''(~1)Gamma'(~1)    3
## 3          alpha(~site)sigma(~1)U(~site)Phi(~1)Gamma''(~1)Gamma'(~1)    4
##       AICc DeltaAICc       weight Deviance
## 4 855.3908  0.000000 5.535834e-01 845.2658
## 2 857.1925  1.801710 2.248779e-01 849.1093
## 5 857.2225  1.831715 2.215293e-01 845.0471
## 1 878.3273 22.936543 5.788635e-06 872.2775
## 3 879.2648 23.874000 3.622512e-06 871.1816
```

```r
best_model <- get(paste0("mabuya_model_", which(tbDT[,modelPosition == 1])))
```

##### Step 9: Interpreting Best Model Results

```r
betaTable <- data.table(best_model$results$beta, keep.rownames = TRUE)
names(betaTable)[names(betaTable) == "rn"] <- "parameter"
betaTable[, Z.value := estimate / se]
betaTable[, p.value := 2 * (1 - pnorm(abs(Z.value)))]

print("Parameter estimates from best CMR model:")
```

```
## [1] "Parameter estimates from best CMR model:"
```

```r
print(betaTable)
```

```
##            parameter   estimate        se        lcl        ucl    Z.value
##               <char>      <num>     <num>      <num>      <num>      <num>
## 1: alpha:(Intercept) -1.9612543 0.1597802 -2.2744234 -1.6480852 -12.274702
```

```
## 2:      alpha:effort  0.0205160 0.0040357  0.0126060  0.0284260   5.083629
## 3:   alpha:sitePARNA  0.4154279 0.2066882  0.0103191  0.8205367   2.009926
## 4:     U:(Intercept)  4.2427291 0.1249664  3.9977950  4.4876632  33.950959
## 5:       U:sitePARNA -2.0009952 0.2402727 -2.4719298 -1.5300606  -8.328017
##         p.value
##           <num>
## 1: 0.000000e+00
## 2: 3.702916e-07
## 3: 4.443907e-02
## 4: 0.000000e+00
## 5: 0.000000e+00
```

This table shows us that alpha (resighting rate/detection probability) is significantly **HIGHER** in PARNA while U (population size) is significantly **LOWER** in PARNA. So, this means that we DO see/capture more in PARNA, and yet, the population size is considerably smaller. This is due to a clear pressure of invasive species and the potential effects of food supplementation.

**Note**: We did not manage capture-recapture on Secondary islands due to impossible logistics.

### Body Size Analysis

Using head length as the most stable morphometric variable.

#### Step 1: Loading Size Data

```r
mabuia_ds <- fread("data/Size_Data.csv")
```

#### Step 2: Assessing Size Analysis Sampling Balance

```r
pointsCounts_size <- mabuia_ds[, .N, by = "site"]
pointsCounts_size[site == "APA", pointsArea := 5.466]
pointsCounts_size[site == "PARNA", pointsArea := 11.424]
pointsCounts_size[site == "Secondary", pointsArea := 1.33]
pointsCounts_size[, pointsDensity := N/pointsArea]
pointsCounts_size[, propPoints := N/(sum(pointsCounts_size[, N]))]
pointsCounts_size[, propSize := pointsArea/(sum(pointsCounts_size[, pointsArea]))]

print("Size analysis sampling distribution:")
```

```
## [1] "Size analysis sampling distribution:"
```

```r
print(pointsCounts_size)
```

```
## Index: <site>
##         site     N pointsArea pointsDensity propPoints   propSize
##       <char> <int>      <num>         <num>      <num>      <num>
## 1:       APA    58      5.466     10.611050  0.4754098 0.30000000
## 2:     PARNA    44     11.424      3.851541  0.3606557 0.62700329
## 3: Secondary    20      1.330     15.037594  0.1639344 0.07299671
```

There is indeed a bit of unbalance but it is not as critical as it seems:

- **APA**: 47% points over 30% area
- **PARNA**: 36% points over 62% area
- **Secondary**: 16% points over 7% area

##### Step 3: Factor Level Setup

```
mabuia_ds[, site := factor(site, levels = c("Secondary", "APA","PARNA"))] # "Secondary" as reference
mabuia_ds[, sex := factor(sex)] # R will pick a reference, usually "F" alphabetically
```

##### Step 4: Outlier Detection

```
detect_outlier <- function(x) {
  Quantile1 <- quantile(x, probs=.25)
  Quantile3 <- quantile(x, probs=.75)
  IQR = Quantile3-Quantile1
  x > Quantile3 + (IQR*1.5) | x < Quantile1 - (IQR*1.5)
}

outliers <- detect_outlier(mabuia_ds$headLength[!is.na(mabuia_ds$headLength)])
print(paste("Number of outliers detected:", sum(outliers)))
```

```
## [1] "Number of outliers detected: 0"
```

No outliers detected, indicating good data quality.

##### Step 5: Head Length Model

```
Head <- glm(headLength ~ sex + site,
                 family = Gamma(link = "log"),
                 data = mabuia_ds)
summary(Head)
```

```
##
## Call:
## glm(formula = headLength ~ sex + site, family = Gamma(link = "log"),
##     data = mabuia_ds)
##
## Coefficients:
##             Estimate Std. Error t value Pr(>|t|)
## (Intercept)  0.50760    0.03862  13.143  < 2e-16 ***
## sexM         0.16930    0.02902   5.833 5.01e-08 ***
## siteAPA     -0.10267    0.03909  -2.627  0.00979 **
## sitePARNA   -0.09844    0.04095  -2.404  0.01780 *
## ---
## Signif. codes:  0 '***' 0.001 '**' 0.01 '*' 0.05 '.' 0.1 ' ' 1
##
## (Dispersion parameter for Gamma family taken to be 0.02271701)
##
##     Null deviance: 3.7328  on 119  degrees of freedom
## Residual deviance: 2.7855  on 116  degrees of freedom
##   (2 observations deleted due to missingness)
## AIC: 25.051
##
## Number of Fisher Scoring iterations: 4
```

#### Step 6: Model Diagnostics

```r
hist(residuals(Head), main = "Residuals Distribution")
```

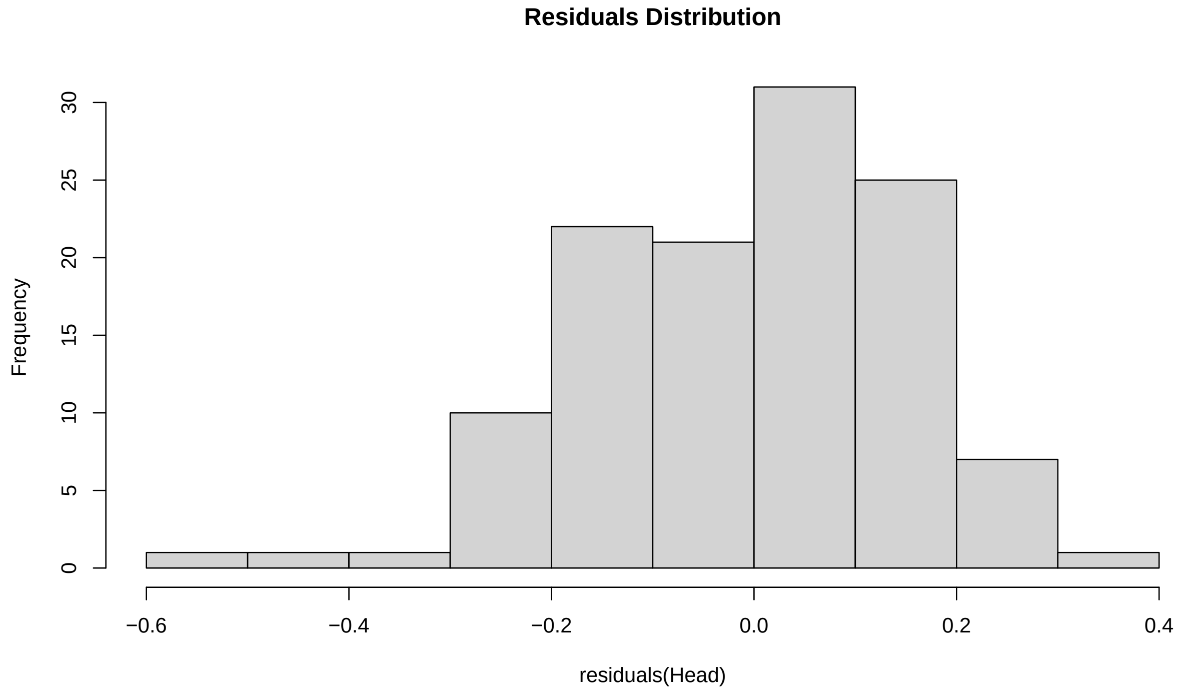

```r
par(mfrow=c(2,2))
plot(Head)
```

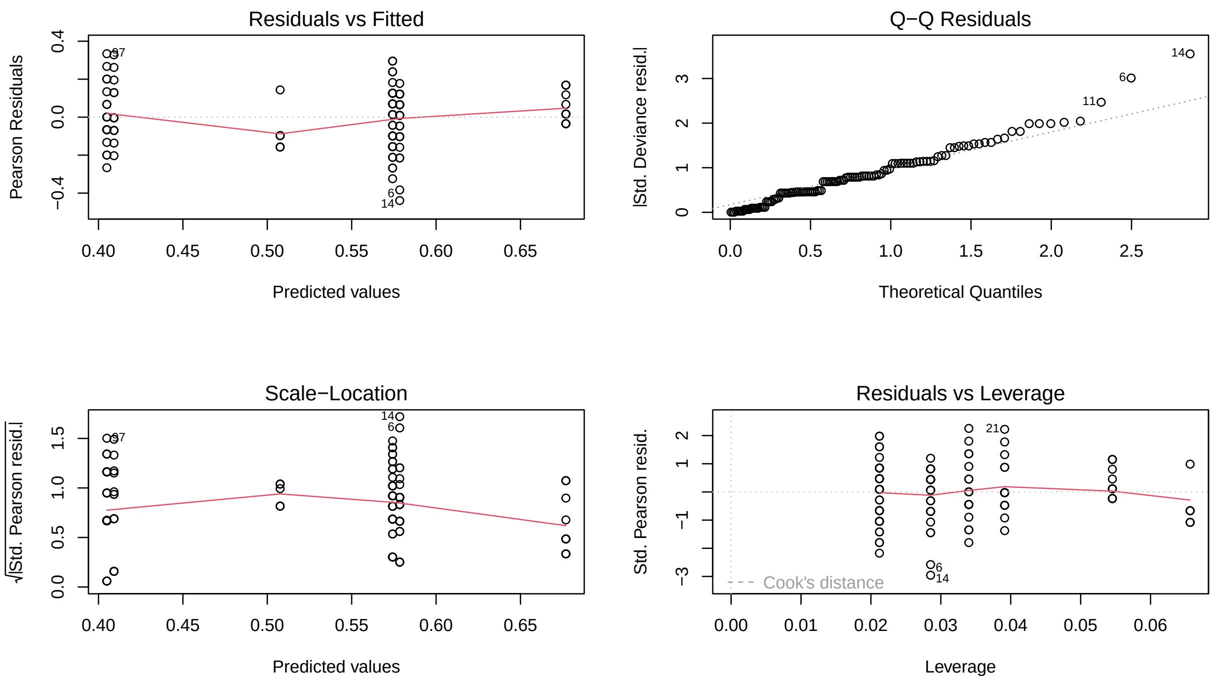

```r
par(mfrow=c(1,1))
```

##### Step 7: Estimated Marginal Means for Head Length

```r
emm_hF <- emmeans(Head, specs = ~ site + sex)
pairwise_head <- pairs(emm_hF, adjust = "tukey")
emm_hDT <- data.table(summary(emm_hF, type = "response"))

print("Head length estimated marginal means:")
```

```
## [1] "Head length estimated marginal means:"
```

```r
print(emm_hDT)
```

```
##         site    sex response         SE    df lower.CL upper.CL
##       <fctr> <fctr>    <num>      <num> <num>    <num>    <num>
## 1: Secondary      F 1.661307 0.06416478   116 1.538960 1.793380
## 2:       APA      F 1.499204 0.04166956   116 1.418903 1.584050
## 3:     PARNA      F 1.505558 0.04488972   116 1.419223 1.597146
## 4: Secondary      M 1.967770 0.06926544   116 1.835254 2.109854
## 5:       APA      M 1.775764 0.03898873   116 1.700197 1.854690
## 6:     PARNA      M 1.783290 0.04540648   116 1.695587 1.875529
```

```r
print("Pairwise comparisons:")
```

```
## [1] "Pairwise comparisons:"
```

```r
print(pairwise_head)
```

```
##  contrast                   estimate     SE  df t.ratio p.value
##  Secondary F - APA F         0.10267 0.0391 116   2.627  0.0991
##  Secondary F - PARNA F       0.09844 0.0409 116   2.404  0.1634
##  Secondary F - Secondary M -0.16930 0.0290 116  -5.833  <.0001
##  Secondary F - APA M       -0.06663 0.0483 116  -1.380  0.7392
##  Secondary F - PARNA M     -0.07086 0.0503 116  -1.408  0.7219
##  APA F - PARNA F           -0.00423 0.0305 116  -0.138  1.0000
##  APA F - Secondary M       -0.27197 0.0491 116  -5.542  <.0001
##  APA F - APA M             -0.16930 0.0290 116  -5.833  <.0001
##  APA F - PARNA M           -0.17353 0.0427 116  -4.061  0.0012
##  PARNA F - Secondary M     -0.26774 0.0501 116  -5.347  <.0001
##  PARNA F - APA M           -0.16507 0.0415 116  -3.973  0.0017
##  PARNA F - PARNA M         -0.16930 0.0290 116  -5.833  <.0001
##  Secondary M - APA M        0.10267 0.0391 116   2.627  0.0991
##  Secondary M - PARNA M      0.09844 0.0409 116   2.404  0.1634
##  APA M - PARNA M           -0.00423 0.0305 116  -0.138  1.0000
##
## Results are given on the log (not the response) scale.
## P value adjustment: tukey method for comparing a family of 6 estimates
```

##### Step 8: Testing for Interaction Effects

```r
Head_Interaction <- glm(headLength ~ sex * site,
                        family = Gamma(link = "log"),
                        data = mabuia_ds)
summary(Head_Interaction)
```

```
##
## Call:
## glm(formula = headLength ~ sex * site, family = Gamma(link = "log"),
##     data = mabuia_ds)
##
## Coefficients:
##                 Estimate Std. Error t value Pr(>|t|)
## (Intercept)     0.414944   0.056381   7.360 3.07e-11 ***
## sexM            0.308509   0.069931   4.412 2.34e-05 ***
## siteAPA        -0.002486   0.065954  -0.038   0.9700
## sitePARNA       0.025459   0.068280   0.373   0.7099
## sexM:siteAPA   -0.150422   0.081438  -1.847   0.0673 .
## sexM:sitePARNA -0.188238   0.084841  -2.219   0.0285 *
## ---
## Signif. codes:  0 '***' 0.001 '**' 0.01 '*' 0.05 '.' 0.1 ' ' 1
##
## (Dispersion parameter for Gamma family taken to be 0.02225138)
##
##     Null deviance: 3.7328  on 119  degrees of freedom
## Residual deviance: 2.6737  on 114  degrees of freedom
##   (2 observations deleted due to missingness)
## AIC: 24.116
##
## Number of Fisher Scoring iterations: 4
```

```
emm_interaction <- emmeans(Head_Interaction, specs = ~ sex * site)
pairs_interaction_head <- pairs(emm_interaction, adjust = "tukey")
print("Interaction model pairwise comparisons:")
```

```
## [1] "Interaction model pairwise comparisons:"
```

```
print(pairs_interaction_head)
```

```
##  contrast                  estimate     SE  df t.ratio p.value
##  F Secondary - M Secondary -0.30851 0.0699 114  -4.412  0.0003
##  F Secondary - F APA        0.00249 0.0660 114   0.038  1.0000
##  F Secondary - M APA       -0.15560 0.0612 114  -2.541  0.1210
##  F Secondary - F PARNA     -0.02546 0.0683 114  -0.373  0.9990
##  F Secondary - M PARNA     -0.14573 0.0633 114  -2.303  0.2013
##  M Secondary - F APA        0.31099 0.0537 114   5.792  <.0001
##  M Secondary - M APA        0.15291 0.0478 114   3.201  0.0214
##  M Secondary - F PARNA      0.28305 0.0565 114   5.008  <.0001
##  M Secondary - M PARNA      0.16278 0.0504 114   3.233  0.0195
##  F APA - M APA             -0.15809 0.0417 114  -3.788  0.0033
##  F APA - F PARNA           -0.02795 0.0515 114  -0.542  0.9943
##  F APA - M PARNA           -0.14822 0.0447 114  -3.318  0.0151
##  M APA - F PARNA            0.13014 0.0453 114   2.872  0.0537
##  M APA - M PARNA            0.00987 0.0373 114   0.264  0.9998
##  F PARNA - M PARNA         -0.12027 0.0480 114  -2.504  0.1317
##
## Results are given on the log (not the response) scale.
## P value adjustment: tukey method for comparing a family of 6 estimates
```

##### Step 9: Ecological Interpretation of Size Patterns

```
dt <- as.data.table(emm_interaction)

# Compare sex ratios across sites
male_props <- table(mabuia_ds[sex == "M", site])/NROW(mabuia_ds[sex == "M",])
female_props <- table(mabuia_ds[sex == "F", site])/NROW(mabuia_ds[sex == "F",])

print("Proportion of males by site:")
```

```
## [1] "Proportion of males by site:"
```

```
print(male_props)
```

```
##
## Secondary       APA     PARNA
## 0.1645570 0.4936709 0.3417722
```

```
print("Proportion of females by site:")
```

```
## [1] "Proportion of females by site:"
```

```
print(female_props)
```

```
##
## Secondary       APA     PARNA
## 0.1627907 0.4418605 0.3953488
```

#### Injury Analysis (Autotomy)

##### Step 1: Creating Autotomy Variable

```
mabuia_ds[, has_autotomy := ifelse(autotomy == "", "No", "Yes")]
```

##### Step 2: Autotomy Proportions and Statistical Test

```
autotomy_table_3way <- table(mabuia_ds$site, mabuia_ds$has_autotomy)
propTable <- prop.table(autotomy_table_3way, margin = 1)
chi_test_autotomy_3way <- chisq.test(autotomy_table_3way)
correctedProportion <- round(propTable*100, 2)
colnames(correctedProportion) <- c("No autotomy signs", "Autotomy signs")

print("Autotomy proportions by site (%):")
```

```
## [1] "Autotomy proportions by site (%):"
```

```
print(correctedProportion)
```

```
##
##             No autotomy signs Autotomy signs
##   Secondary             85.00          15.00
##   APA                   70.69          29.31
##   PARNA                 75.00          25.00
```

```
print("Chi-square test results:")
```

```
## [1] "Chi-square test results:"
```

```
print(chi_test_autotomy_3way)
```

```
##
##  Pearson's Chi-squared test
##
## data:  autotomy_table_3way
## X-squared = 1.613, df = 2, p-value = 0.4464
```

##### Post-hoc power analysis for injury (autotomy)

```
tab <- autotomy_table_3way
expected <- chi_test_autotomy_3way$expected
w <- sqrt(sum((tab - expected)^2 / expected) / sum(tab))  # effect size (Cohen's w)
N <- sum(tab)
df_chisq <- (nrow(tab) - 1) * (ncol(tab) - 1)
pwr_overall <- pwr.chisq.test(w = w, N = N, df = df_chisq, sig.level = 0.05)
pwr_needed <- pwr.chisq.test(w = w, power = 0.80, df = df_chisq, sig.level = 0.05)

cat("Omnibus 3×2 power (observed effect):", round(pwr_overall$power, 3), "\n")
```

```
## Omnibus 3×2 power (observed effect): 0.189
```

```
cat("Total N required for 80% power (observed effect):", ceiling(pwr_needed$N), "\n")
```

```
## Total N required for 80% power (observed effect): 729
```

```
cat("Cohen's w:", round(w, 3), "with N =", N, "and df =", df_chisq, "\n\n")
```

```
## Cohen's w: 0.115 with N = 122 and df = 2
```

```
# 3) Pairwise power (two-proportion tests) with actual n's
yes <- tab[, "Yes"]
no  <- tab[, setdiff(colnames(tab), "Yes")]
n   <- yes + no
p   <- yes / n
site_levels <- rownames(tab)

# helper for unequal-n power using Cohen's h
pair_power <- function(p1, p2, n1, n2, alpha = 0.05) {
  ES_h <- 2*asin(sqrt(p1)) - 2*asin(sqrt(p2))
  out <- pwr.2p2n.test(h = abs(ES_h), n1 = n1, n2 = n2, sig.level = alpha)
  out$power
}

pairs <- t(combn(seq_along(site_levels), 2))
pw_out <- lapply(seq_len(nrow(pairs)), function(k){
  i <- pairs[k,1]; j <- pairs[k,2]
  data.frame(
    contrast = paste(site_levels[i], "vs", site_levels[j]),
    n1 = as.integer(n[i]), n2 = as.integer(n[j]),
    p1 = round(p[i],3), p2 = round(p[j],3),
    power_0.05 = round(pair_power(p[i], p[j], n[i], n[j], 0.05), 3)
  )
})
pairwise_power_table <- do.call(rbind, pw_out)
print(pairwise_power_table, row.names = FALSE)
```

```
##            contrast n1 n2    p1    p2 power_0.05
##    Secondary vs APA 20 58 0.150 0.293      0.270
##  Secondary vs PARNA 20 44 0.150 0.250      0.154
##        APA vs PARNA 58 44 0.293 0.250      0.077
```

```r
# 4) Minimum Detectable Difference (MDD) for key contrasts at 80% power
mdd_for_pair <- function(n1, n2, p_baseline, power_target = 0.80, alpha = 0.05) {
  deltas <- seq(0.001, 0.40, by = 0.001)
  pow <- sapply(deltas, function(d){
    p1 <- p_baseline
    p2 <- max(min(p_baseline + d, 0.99), 0.01)
    ES_h <- 2*asin(sqrt(p1)) - 2*asin(sqrt(p2))
    pwr.2p2n.test(h = abs(ES_h), n1 = n1, n2 = n2, sig.level = alpha)$power
  })
  idx <- which(pow >= power_target)
  if (length(idx) == 0) return(NA_real_)
  deltas[min(idx)]
}

# Choose contrasts that involve the smallest sample (often 'Secondary')
sec_ix  <- which(site_levels == "Secondary")
apa_ix  <- which(site_levels == "APA")
par_ix  <- which(site_levels == "PARNA")

mdd_list <- list()
if (length(sec_ix) && length(apa_ix)) {
  mdd_list[["MDD_Sec_vs_APA"]] <- round(100*mdd_for_pair(n[sec_ix], n[apa_ix], p[sec_ix], 0.80), 1)
}
if (length(sec_ix) && length(par_ix)) {
  mdd_list[["MDD_Sec_vs_PARNA"]] <- round(100*mdd_for_pair(n[sec_ix], n[par_ix], p[sec_ix], 0.80), 1)
}
print(mdd_list)
```

```
## $MDD_Sec_vs_APA
## [1] 32.6
##
## $MDD_Sec_vs_PARNA
## [1] 34.1
```

```r
# 5) Simulation-based omnibus power under observed prevalences
set.seed(123)
sim_power_chisq <- function(n_by_site, p_by_site, B = 5000, alpha = 0.05) {
  stopifnot(length(n_by_site) == length(p_by_site))
  hits <- 0L
  for (b in seq_len(B)) {
    sim_yes <- rbinom(length(n_by_site), size = n_by_site, prob = p_by_site)
    sim_no  <- n_by_site - sim_yes
    sim_tab <- cbind(sim_yes, sim_no)
    dimnames(sim_tab) <- list(names(n_by_site), c("Yes","No"))
    pval <- suppressWarnings(chisq.test(sim_tab, correct = FALSE)$p.value)
    if (pval < alpha) hits <- hits + 1L
  }
  hits / B
}
```

```r
n_by_site <- setNames(as.numeric(n), site_levels)
p_by_site <- setNames(as.numeric(p), site_levels)
sim_power <- sim_power_chisq(n_by_site, p_by_site, B = 5000, alpha = 0.05)
cat("Simulation-based omnibus power:", round(sim_power, 3), "\n")
```

```
## Simulation-based omnibus power: 0.16
```

> *Reporting note:* Using the observed 3×2 table, Cohen's $w = 0.115$. With our sample size (122), the omnibus test has power approximately 0.19 to detect this effect; achieving 80% power would require ~729 total observations. Pairwise power was lowest for contrasts involving the smallest sample, explaining the non-significant test despite consistent directional patterns. With current n, the minimum detectable absolute difference for comparisons involving the secondary islands is 32.6–34.1 percentage points to reach 80% power.

Autotomy was proportionally more frequent in the APA than in PARNA and roughly twice that observed on the secondary islands, suggesting elevated predation pressure on the main island. However, these differences were not statistically significant. Our power analyses indicate that this outcome is largely due to limited sample sizes—particularly on the secondary islands—rather than the absence of a biological effect, highlighting the need for expanded sampling to better resolve these patterns.

### Combined Size and Injury Visualization

#### Step 1: Preparing Interaction Data for Plotting

```r
dt_df <- as.data.frame(dt)
dt_df$site <- factor(dt_df$site, levels = c("Secondary", "APA", "PARNA"))

# Get significance testing for interaction model
pairs_interaction <- contrast(emm_interaction, method = "pairwise", adjust = "tukey")
summary_pw_int <- summary(pairs_interaction)
p_values_int <- summary_pw_int$p.value
print(p_values_int)
```

```
##  [1] 3.312743e-04 1.000000e+00 1.210025e-01 9.990378e-01 2.012888e-01
##  [6] 9.272989e-07 2.142024e-02 2.956024e-05 1.949409e-02 3.262026e-03
## [11] 9.942631e-01 1.505577e-02 5.373153e-02 9.998207e-01 1.316787e-01
```

#### Step 2: Processing Significance Letters for Interaction

```r
contrast_names_raw_int <- as.character(summary_pw_int$contrast)
p_value_names <- sapply(contrast_names_raw_int, function(contrast) {
  parts <- strsplit(contrast, " - ")[[1]]
  key1 <- gsub(" ", "", parts[1])
  key2 <- gsub(" ", "", parts[2])
  return(paste(key1, key2, sep = "-"))
})
names(p_values_int) <- p_value_names

letters_result_int <- multcompLetters(p_values_int)
significance_letters_int <- letters_result_int$Letters

dt_df$key <- paste0(dt_df$sex, dt_df$site)
dt_df$significance_group <- significance_letters_int[dt_df$key]
```

```r
dt_df$y_pos_letters <- dt_df$upper.CL + 0.03
plot(dt_df)
```

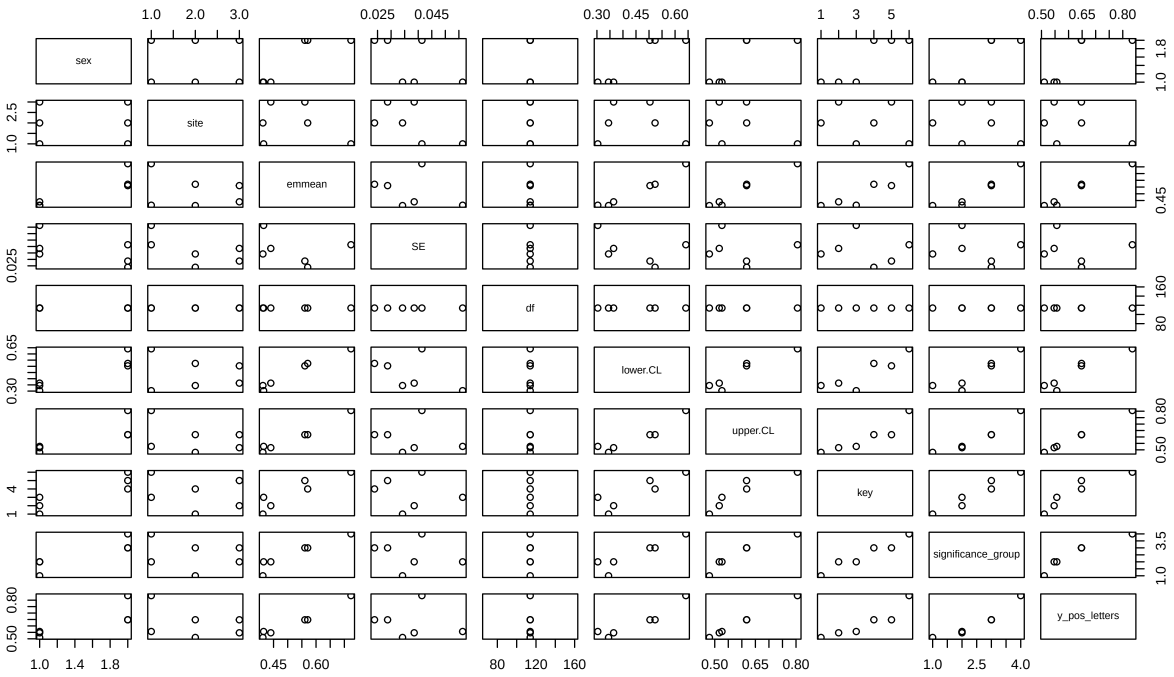

##### Step 3: Preparing Autotomy Data by Sex

```r
# For males
males_data <- mabuia_ds[mabuia_ds$sex == 'M', ]
prop_table_males <- prop.table(table(males_data$site, males_data$has_autotomy), margin = 1)
males_autotomy_df <- as.data.frame.matrix(prop_table_males)
names(males_autotomy_df)[names(males_autotomy_df) == "Yes"] <- "injury_prop"
males_autotomy_df$site <- rownames(males_autotomy_df)
males_autotomy_df$sex <- "Males"

# For females
females_data <- mabuia_ds[mabuia_ds$sex == 'F', ]
prop_table_females <- prop.table(table(females_data$site, females_data$has_autotomy), margin = 1)
females_autotomy_df <- as.data.frame.matrix(prop_table_females)
names(females_autotomy_df)[names(females_autotomy_df) == "Yes"] <- "injury_prop"
females_autotomy_df$site <- rownames(females_autotomy_df)
females_autotomy_df$sex <- "Females"

# Combine
autotomy_df_final <- rbind(males_autotomy_df, females_autotomy_df)
autotomy_df_final$site <- factor(autotomy_df_final$site, levels = c("Secondary", "APA", "PARNA"))
autotomy_df_final$sex <- factor(autotomy_df_final$sex, levels = c("Females", "Males"))
plot(autotomy_df_final)
```

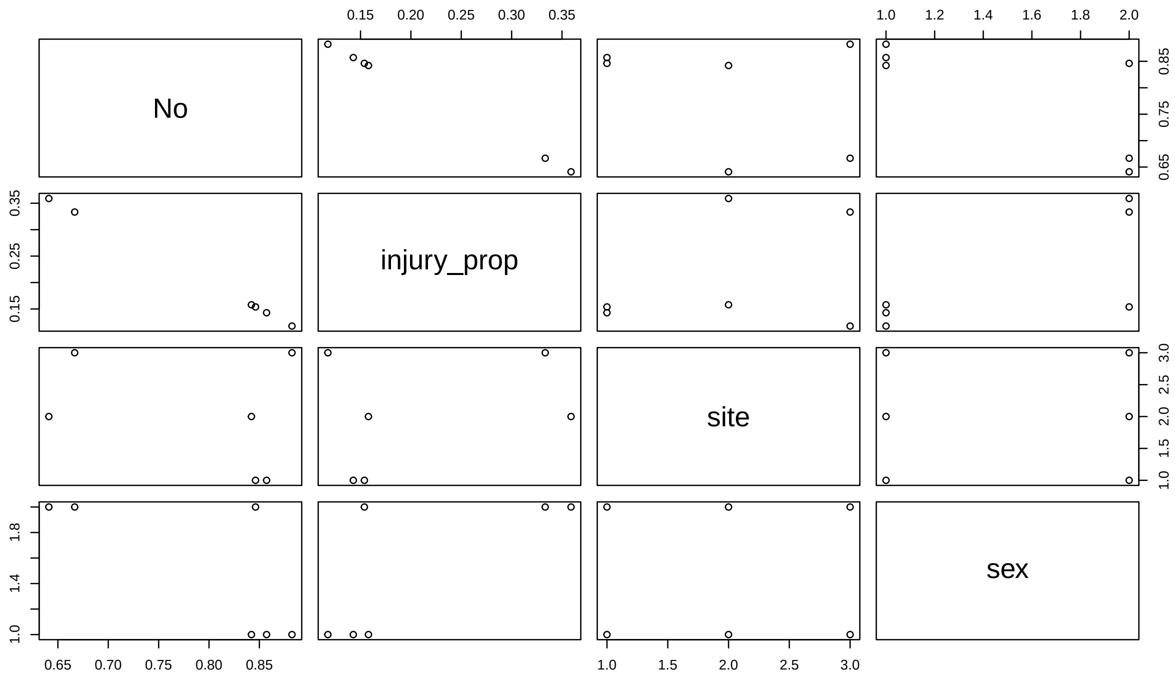

##### Step 4: Formatting for Unified Plot

```
dt_df$sex <- ifelse(dt_df$sex == "F", "Females", "Males")
dt_df$sex <- factor(dt_df$sex, levels = c("Females", "Males"))

# Create unified plotting dataset
# Select and rename columns from head length data
plot_data_head <- data.frame(
  site = dt_df$site,
  sex = dt_df$sex,
  measurement = "Head Length (cm)",
  value = dt_df$emmean,
  lower_ci = dt_df$lower.CL,
  upper_ci = dt_df$upper.CL,
  significance_group = dt_df$significance_group
)

# Select and rename columns from injury data
plot_data_injury <- data.frame(
  site = autotomy_df_final$site,
  sex = autotomy_df_final$sex,
  measurement = "Proportion Injury",
  value = autotomy_df_final$injury_prop,
  lower_ci = NA, # No error bars for injury rate
  upper_ci = NA,
  significance_group = NA # No significance letters for injury rate
)

plot_data_unified <- rbind(plot_data_head, plot_data_injury)
```

```r
plot(plot_data_unified)
```

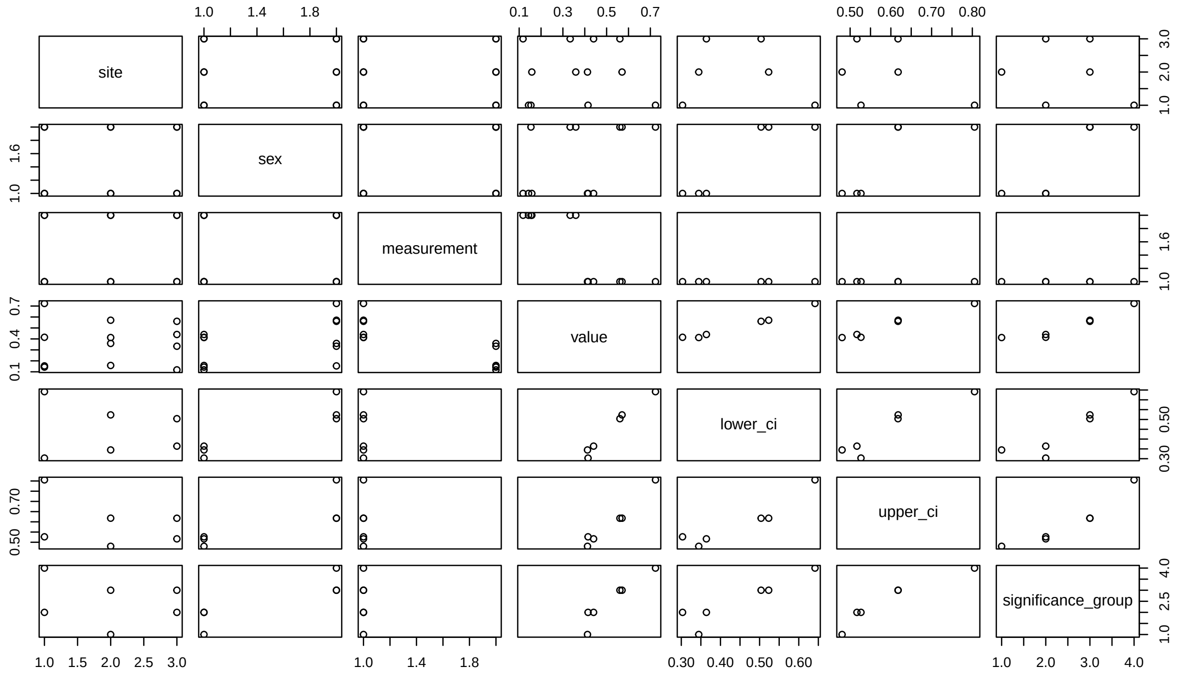

##### Creating the Final Size and Injury Plot

```r
body_size_plot <- ggplot(
  plot_data_unified,
  aes(x = site, y = value, fill = site, color = site)
) +
    # --- Geoms that use SUBSETS of the data ---
  # Add bars ONLY for the injury rate data
  geom_bar(
    data = plot_data_unified[plot_data_unified$measurement == "Proportion Injury", ],
    stat = "identity", color = "black", alpha = 0.8
  ) +
  # Add error bars ONLY for the head length data
  geom_errorbar(
    data = plot_data_unified[plot_data_unified$measurement == "Head Length (cm)", ],
    aes(ymin = lower_ci, ymax = upper_ci), width = 0.25, linewidth = 1
  ) +
  # Add points ONLY for the head length data
  geom_point(
    data = plot_data_unified[plot_data_unified$measurement == "Head Length (cm)", ],
    size = 4, stroke = 1.5
  ) +
  # Add text ONLY for the head length data
  geom_text(
    data = plot_data_unified[plot_data_unified$measurement == "Head Length (cm)", ],
    aes(y = upper_ci + 0.1, label = significance_group), # Increase offset for % scale
    color = "black", size = 6, fontface = "bold"
```

```r
) +

# --- Faceting ---
# Facet by both sex and measurement, and allow y-axes to be independent
facet_grid(
  measurement ~ sex,
  scales = "free_y"
) +

# --- Scales and Labels ---
scale_color_viridis_d(option = "D") +
scale_fill_viridis_d(option = "D") +

# --- Theming ---
theme_bw(base_size = 14) +
theme(
  legend.position = "none",
  plot.title = element_text(face = "bold", size = 16),
  strip.text = element_text(size = 12, face = "bold"),
  strip.placement = "outside", # Ensures strips are next to the axis
  axis.title = element_text(face = "bold"),
  panel.grid.major.x = element_blank(),
  panel.grid.minor = element_blank(),
  # Add a bit more space between facets
  panel.spacing = unit(1.0, "lines"),
  axis.title.x = element_blank(),
  axis.title.y = element_blank(),
)
```

#### Summarizing the findings:

##### Results from point counts:

```r
summary(emm_density_hierarchical, infer = TRUE)
```

```
##  InvasiveSpeciesPresence FoodSupplementation response     SE  df asymp.LCL
##  0                       0                      0.411 0.0696 Inf     0.295
##  1                       0                      0.156 0.0247 Inf     0.115
##  0                       1                      0.537 0.1476 Inf     0.314
##  1                       1                      0.204 0.0301 Inf     0.153
##  asymp.UCL null z.ratio p.value
##      0.573    1  -5.252  <.0001
##      0.213    1 -11.730  <.0001
##      0.921    1  -2.261  0.0238
##      0.273    1 -10.782  <.0001
##
## Confidence level used: 0.95
## Intervals are back-transformed from the log scale
## Tests are performed on the log scale
```

```r
density_plot
```

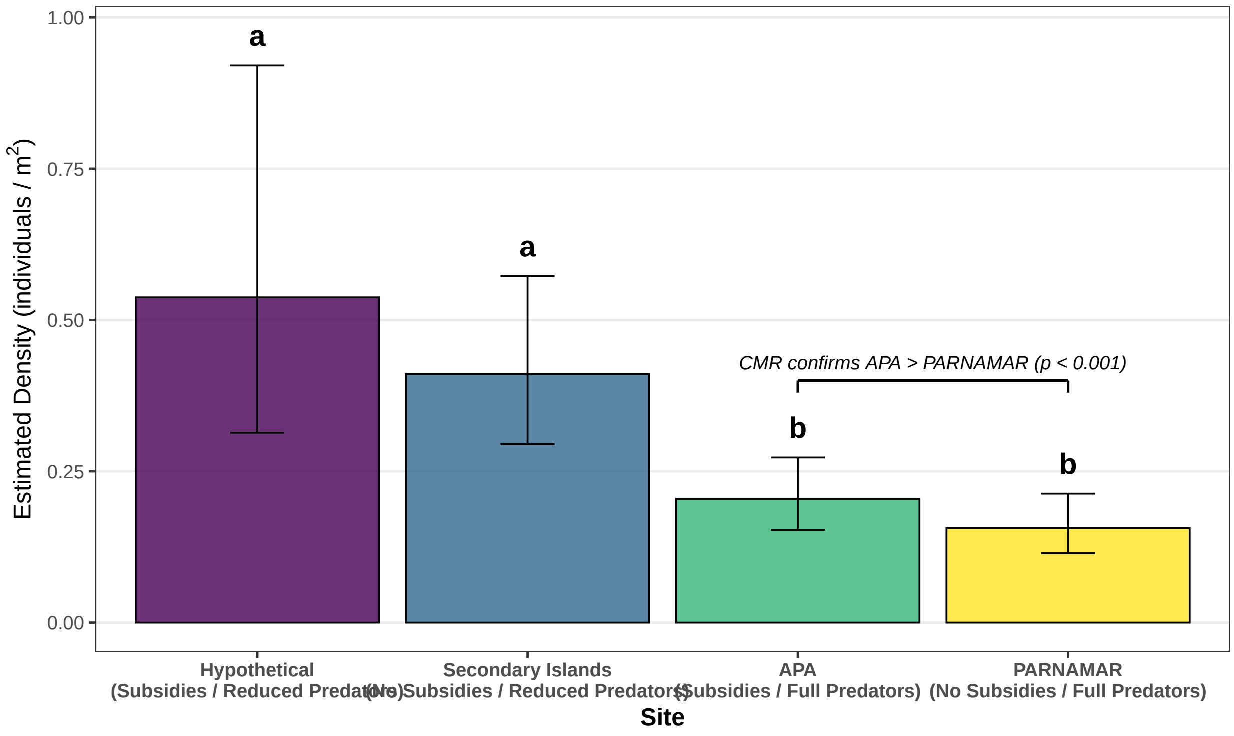

#### Results from CMR:

```r
print(betaTable)
```

```
##             parameter   estimate        se        lcl        ucl    Z.value
##                <char>      <num>     <num>      <num>      <num>      <num>
## 1: alpha:(Intercept) -1.9612543 0.1597802 -2.2744234 -1.6480852 -12.274702
## 2:       alpha:effort  0.0205160 0.0040357  0.0126060  0.0284260   5.083629
## 3:    alpha:sitePARNA  0.4154279 0.2066882  0.0103191  0.8205367   2.009926
## 4:     U:(Intercept)  4.2427291 0.1249664  3.9977950  4.4876632  33.950959
## 5:       U:sitePARNA -2.0009952 0.2402727 -2.4719298 -1.5300606  -8.328017
##          p.value
##            <num>
## 1: 0.000000e+00
## 2: 3.702916e-07
## 3: 4.443907e-02
## 4: 0.000000e+00
## 5: 0.000000e+00
```

#### Results from size analysis:

```r
body_size_plot
```

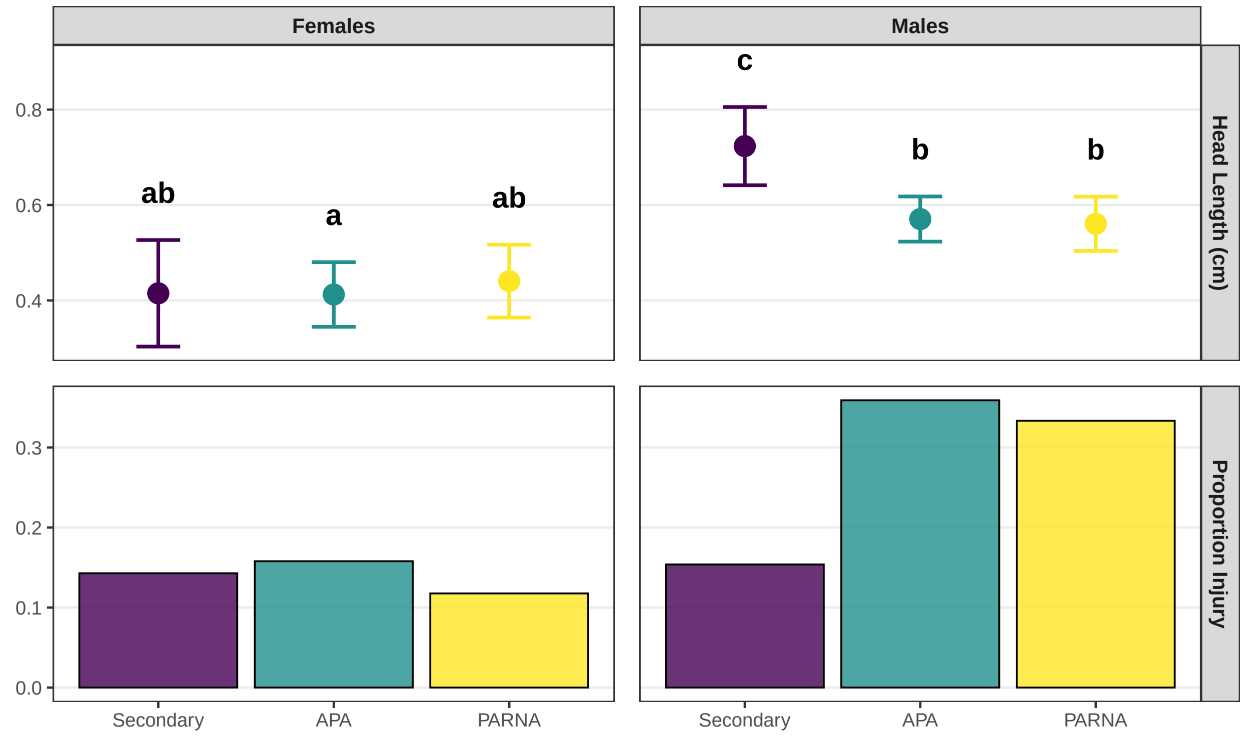

#### Results from sublethal injuries:

```r
print(correctedProportion)
```

```
##
##               No autotomy signs Autotomy signs
##   Secondary               85.00          15.00
##   APA                     70.69          29.31
##   PARNA                   75.00          25.00
```

```r
print(chi_test_autotomy_3way)
```

```
##
##  Pearson's Chi-squared test
##
## data:  autotomy_table_3way
## X-squared = 1.613, df = 2, p-value = 0.4464
```

#### Saving the plots for manuscript

```r
ggsave(
  filename = "outputs/Density_Plot.png", # The name of the output file
  plot = density_plot,                     # The plot object to save
  width = 12,                               # Width of the image in inches
  height = 6,                              # Height of the image in inches
  dpi = 300,                               # Dots per inch (standard for publication)
  bg = "white"                             # Set a white background
)
```

```r
# Save as a high-resolution TIFF file
ggsave(
  filename = "outputs/Density_Plot.tiff",
  plot = density_plot,
  width = 12,
  height = 6,
  dpi = 300
)

ggsave(
  filename = "outputs/Size_Injury.png",
  plot = body_size_plot,
  width = 8,   # Keep it reasonably wide for the two columns
  height = 7,  # A bit taller to accommodate the stacked facets
  dpi = 300,
  bg = "white"
)

ggsave(
  filename = "outputs/Size_Injury.tiff",
  plot = body_size_plot,
  width = 8,   # Keep it reasonably wide for the two columns
  height = 7,  # A bit taller to accommodate the stacked facets
  dpi = 300,
  bg = "white"
)
```
